# Whole genome comparisons of ergot fungi reveals the divergence and evolution of species within the genus *Claviceps* are the result of varying mechanisms driving genome evolution and host range expansion

**DOI:** 10.1101/2020.04.13.039230

**Authors:** Stephen A. Wyka, Stephen J. Mondo, Miao Liu, Jeremy Dettman, Vamsi Nalam, Kirk D. Broders

## Abstract

The genus *Claviceps* has been known for centuries as an economically important fungal genera for pharmacology and agricultural research. Only recently have researchers begun to unravel the evolutionary history of the genus, with origins in South America and classification of four distinct sections through ecological, morphological, and metabolic features (*Claviceps* sects. *Citrinae, Paspalorum*, *Pusillae*, and *Claviceps*). The first three sections are additionally characterized by narrow host range, while sect. *Claviceps* is considered evolutionarily more successful and adaptable as it has the largest host range and biogeographical distribution. However, the reasons for this success and adaptability remain unclear. Our study elucidates factors influencing adaptability by sequencing and annotating 50 *Claviceps* genomes, representing 21 species, for a comprehensive comparison of genome architecture and plasticity in relation to host range potential. Our results show the trajectory from specialized one-speed genomes (sects. *Citrinae* and *Paspalorum*) towards adaptive two-speed genomes (sects. *Pusillae* and *Claviceps*) through co-localization of transposable elements around predicted effectors and a putative loss of repeat-induced point mutation resulting in unconstrained tandem gene duplication coinciding with increased host range potential and speciation. Alterations of genomic architecture and plasticity can substantially influence and shape the evolutionary trajectory of fungal pathogens and their adaptability. Furthermore, our study provides a large increase in available genomic resources to propel future studies of *Claviceps* in pharmacology and agricultural research, as well as, research into deeper understanding of the evolution of adaptable plant pathogens.

## Introduction

Fungi, particularly phytopathogenic species, are increasingly being utilized to gain insight into the evolution of eukaryotic organisms, due to their adaptive nature and unique genome structures (Gladieux *et al.* 2014; Dong *et al.* 2015). Adaptation and diversification of fungal species can be mediated by changes in genome architecture and plasticity such as genome size, transposable element (TE) content, localization of TEs to specific genes, genome compartmentalization, gene duplication rates, recombination rates, and presence/absence polymorphism of virulence factors (Dong *et al.* 2015; Möller & Stukenbrock 2017). The presence or absence of repeat-induced point (RIP) mutation is also an important mechanism for fungal genome evolution, as RIP works on a genome-wide scale to silence transposable elements and duplicated genes, which can also “leak” onto neighboring genes (Galagan *et al.* 2003, 2004; Raffaele & Kamoun 2012; Urguhart *et al.* 2018; Möller & Stukenbrock 2019). It is becoming increasingly evident that variations in these factors can be used to classify genomes as a one-speed (one-compartment), such as the powdery mildew fungi *Blumeria graminis* f.sp. *hordei* and f.sp *tritici*, two-speed (two-compartment), such as the late blight pathogen *Phytophthora infestans*, or multi-speed (multi-compartment) such as the multi-host pathogen *Fusarium oxysporum* (Dong *et al.* 2015; Frantzeskakis *et al.* 2019). These different “speeds” are characterized by their potential adaptability such that one-speed genomes are often considered less-adaptable, while two-speed and multi-speed genomes are often considered more-adaptable (Dong *et al.* 2015; Frantzeskakis *et al.* 2019; Möller & Stukenbrock 2019).

The ergot fungi of the genus *Claviceps* (Ascomycota, Hypocreales) are biotrophic species that share a specialized ovarian-specific non-systemic parasitic lifestyle with their grass hosts (Píchová *et al.* 2018). Infections are fully restricted to individual unpollinated ovaries (Tudzynski and Scheffer 2004), and the fungus actively manages to maintain host cell viability to obtain nutrients from living tissue through a complex cross-talk of genes related to pathogenesis, such as secreted effectors, secondary metabolites, or cytokinin production (Hinsch *et al.* 2015, 2016; Oeser *et al.* 2017; Kind *et al.* 2018a, 2018b). Species of *Claviceps* are most notably known for their production of toxic alkaloids and secondary metabolites but are also known for their expansive host range and negative impact on global cereal crop production and livestock farming. These negative effects on human and livestock health are the primary reason *Claviceps* species are referred to as plant pathogens. However, under the light of co-evolution with their grass hosts some *Claviceps* species are considered conditional defensive mutualists with their hosts as they prevent herbivory and can improve host fitness (Raybould *et al.* 1998; Fisher *et al.* 2007; Wäli *et al.* 2013).

The genus *Claviceps* contains 59 species divided into four sections; sects. *Claviceps*, *Pusillae*, *Citrinae*, and *Paspalorum* (Píchová *et al.* 2018). It was postulated that sects. *Citrinae* and *Paspalorum* originated in South America, while sect. *Pusillae* experienced speciation throughout the Eocene, Oligocene, and Miocene as these species encountered newly emergent PACMAD warm-season grasses (subfamilies Panicoideae, Aristidoideae, Chloridoideae, Micrairoideae, Arundinoideae, Danthonioideae) when an ancestral strain was transferred from South America to Africa (Píchová *et al.* 2018). In contrast, the crown node of sect. *Claviceps* is estimated at 20.4 Mya and was followed by a radiation of the section corresponding to a host jump from ancestral sedges (Cyperaceae) to the BOP clade (cool-season grasses; subfamilies Bambusoideae, Oryzoideae (syn: Ehrhartoideae) (Soreng *et al.* 2015), Pooideae) in North America (Bouchenak-Khelladi *et al.* 2010; Píchová *et al.* 2018). Section *Claviceps* has the largest host range with *C. purpurea sensu stricto* (s.s.) having been reported on up to 400 different species in clade BOP (Alderman *et al.* 2004, Píchová *et al.* 2018) across six tribes, and retains the ability to infect sedges (Cyperaceae) (Jungehülsing and Tudzynski 1997). In contrast, sect. *Pusillae* is specialized to the tribes Paniceae and Andropogoneae, and sects. *Citrinae* and *Paspalorum* only infect members of tribe Paspaleae and tribe Cynodonteae, respectively (Píchová *et al.* 2018). The shared specialized infection life cycle of the *Claviceps* genus, the drastic differences in host range potential of different species, and geographic distribution represent a unique system to study the evolution and host adaptation of eukaryotic organisms.

Despite their ecological and agriculture importance, little is known about the evolution and genomic architecture of these important fungal species in comparison to other cereal pathogens such as species in the genera *Puccinia* (Cantu *et al.* 2013; Kiran *et al.* 2016, 2017), *Zymoseptoria* (Grandaubert *et al.* 2015, 2019; Estep *et al.* 2015; Poppe *et al.* 2015; Testa *et al.* 2015; Wu *et al.* 2017; Stukenbrock & Dutheil 2018), or *Fusarium* (Kvas *et al.* 2009; Ma *et al.* 2010; Rep & Kistler 2010; Watanabe *et al.* 2011; Sperschneider *et al.* 2015). Unfortunately, the lack of genome data for the *Claviceps* genus has hampered our ability to complete comparative analyses to identify factors that are influencing the adaptation of *Claviceps* species across the four sections in the genus, and the mechanisms by which species of sect. *Claviceps* have adapted to such a broad host range, in comparison to the other three sections. Here we present the sequences and annotations of 50 *Claviceps* genomes, representing 19 species, for a comprehensive comparison of the genus to understand evolution within the genus *Claviceps* by characterizing the genomic plasticity and architecture in relation to adaptive host potential. Our analysis reveals the trajectory from specialized one-speed genomes (sects. *Citrinae* and *Paspalorum*) towards adaptive two-speed genomes (sects. *Pusillae* and *Claviceps*) through co-localization of transposable elements around predicted effectors and a putative loss of RIP resulting in tandem gene duplication coinciding with increased host range potential.

## Materials and Methods

### Sample acquisition

Field collected samples (Clav) were surfaced sterilized, allowed to grow as mycelia, and individual conidia transferred to make single spore cultures. Thirteen cultures were provided by Dr. Miroslav Kolařík from the Culture Collection of Clavicipitaceae (CCC) at Institute of Microbiology, Academy of Sciences of the Czech Republic. Raw Illumina reads for samples (LM28, LM582, LM78, LM81, LM458, LM218, LM454, LM576, and LM583) were downloaded from NCBI’s SRA database. Raw Illumina reads from an additional 21 LM samples were generated by Dr. Liu’s lab (AAFC), sequencing protocol of these 21 samples followed Wingfield *et al.* (2018). Summarized information can be found in Additional file 1: Table S1.

### Preparation of genomic DNA

Cultures grown on cellophane PDA plates were used for genomic DNA extraction from lyophilized mycelium following a modified CTAB method (Doyle and Doyle 1987; Wingfield *et al.* 2018) without using the RNase Cocktail™ Enzyme Mix, only RNASE A was used. DNA contamination was checked by running samples on a 1% agarose gel and a NanoDrop One^c^ (Thermo Fishcer Scientific). Twenty samples (7 Clav and 13 CCC) were sent to BGI-Hong Kong HGS Lab for 150-bp paired-end Illumina sequencing on a HiSeq™ 4000.

### Genome assembly

Preliminary data showed that raw reads of LM458 were contaminated with bacterial DNA but showed strong species similarity to Clav32 and Clav50. To filter out the bacterial DNA sequences, reads of LM458 were mapped against the assembled Clav32 and Clav50 genomes using BBSplit v38.41 (Bushnell 2014). All forward and reverse reads mapped to each of the genomes were concatenated, respectively. Both sets were then interleaved to remove duplicates and used for further analysis. Reads for all 50 samples were checked for quality with FastQC v0.11.5 (Andrews 2010) and trimmed with Trimmomatic v0.36 (Bolger *et al.* 2014) using the commands (SLIDINGWINDOW:4:20 MINLEN:36 HEADCROP:10) to remove poor quality data, only paired end reads were used. To better standardize the comparative analysis all 50 sample were subject to *de novo* genome assembly with Shovill v0.9.0 (https://github.com/tseemann/shovill) using SPAdes v3.11.1 (Nurk *et al.* 2013) with a minimum contig length of 1000 bp.

The reference genomes of *C. purpurea* strain 20.1 (SAMEA2272775), *C. fusiformis* PRL 1980 (SAMN02981339), and *C. paspali* RRC 1481 (SAMN02981342) were downloaded from NCBI. Proteins for *C. fusiformis* and *C. paspali* were not available on NCBI so they were extracted from GFF3 files provided by Dr. Chris Schardl and Dr. Neil Moore, University of Kentucky, corresponding to the 2013 annotations (Schardl *et al.* 2013) available at http://www.endophyte.uky.edu. Reference genomes were standardized for comparative analysis with our 50 annotated genomes, by implementing a protein length cutoff of 50 aa and removal of alternatively spliced proteins in *C. fusiformis* and *C. paspali*, only the longest spliced protein for each locus remained.

### Transposable elements

Transposable elements (TE) fragments were identified following procedures for establishment of *de novo* comprehensive repeat libraries set forth in Berriman *et al.* (2018) through a combined use of RepeatModeler v1.0.8 (Smit & Hubley 2015), TransposonPSI (Hass 2010), LTR_finder v1.07 (Xu & Wang 2007), LTR_harvest v1.5.10 (Ellinghaus *et al.* 2008), LTR_digest v1.5.10 (Steinbiss *et al.* 2009), Usearch v11.0.667 (Edgar 2010), and RepeatClassifier v1.0.8 (Smit & Hubley 2015) with the addition of all curated fungal TEs from RepBase (Bao *et al.* 2015). RepeatMasker v4.0.7 (Smit *et al.* 2015) was then used to soft mask the genomes and identify TE regions. TE content was represented as the proportion of the genome masked by TE regions determined by RepeatMasker, excluding simple and low complexity repeats. These steps were automated through construction of a custom script, TransposableELMT (https://github.com/PlantDr430/TransposableELMT).

Divergence landscapes for TEs in all 53 *Claviceps* genomes were generated using a custom script (https://github.com/PlantDr430/CSU_scripts/blob/master/TE_divergence_landscape.py) and the RepeatMasker output results. The RepeatMasker results were also used with the respective GFF3 file from each genome to calculate the average distance (kbp) of each gene to the closest TE fragment on the 5’ and 3’ flanking side. Values were calculated for predicted effectors, non-effector secreted genes, non-secreted metabolite genes, and all other genes using a custom script (https://github.com/PlantDr430/CSU_scripts/blob/master/TE_closeness.py).

### Genome annotation

AUGUSTUS v3.2.2 (Mario *et al.* 2008) was used to create pre-trained parameters files using the reference *C. purpurea* strain 20.1, available EST data from NCBI, and wild-type RNAseq data (SRR4428945) created in Oeser *et al.* (2017). RNA-seq data was subject to quality check and trimming as above. All three datasets were also used to train parameter files for the *ab initio* gene model prediction software’s GeneID v1.4.4 (Blanco *et al.* 2007) and CodingQuarry v2.0 (Testa *et al.* 2015). GeneID training followed protocols available at http://genome.crg.es/software/geneid/training.html. For CodingQuarry training, RNA transcripts were created *de novo* using Trinity v2.8.4 (Grabherr *et al.* 2011) on default settings and EST coordinates were found by mapping the EST data to the reference genome using Minimap2 v2.1 (Li 2018).

Gene models for the 50 genomes were then predicted with GeneID and CodingQuarry using the trained *C. purpruea* parameter files. CodingQuarry prediction was also supplemented with transcript evidence by mapping the available EST and RNA-seq *C. purpurea* data to each genome using Minimap2. BUSCO v3 (Waterhouse *et al.* 2018) was run on all 50 genomes using the AUGUSTUS *C. purpurea* pre-trained parameter files as the reference organism and the Sordariomyceta database. The resulting predicted proteins for each sample were used as training models for *ab initio* gene prediction using SNAP (Korf 2004) and GlimmerHMM v3.0.1 (Majoros *et al.* 2004). Lastly, GeMoMa v1.5.3 (Keilwagen *et al.* 2016) was used for *ab initio* gene prediction using the soft-masked genomes and the *C. purpruea* 20.1 reference files.

Funannotate v1.6.0 (Palmer & Stajich 2019) was then used as the primary software for genome annotation. Funannotate additionally uses AUGUSTUS and GeneMark-ES (Ter-Hovhannisyan *et al.* 2008) for *ab initio* gene model prediction, Exonerate for transcript and protein evidence alignment, and EVidenceModeler (Hass *et al.* 2008) for a final weighted consensus. All *C. purpurea* EST and RNAseq data were used as transcript evidence and the Uniport Swiss-Prot database and proteins from several closely related species (*C. purpurea* strain 20.1, *C. fusiformis* PRL1980, *C. paspali* RRC1481, *Fusarium oxysporum f.sp. lycopersici* 4287, *Pochonia chlamydosporia* 170, *Ustilago maydis* 521, and *Epichloe festucae* F1) were used as protein evidence. The AUGUSTUS pre-trained *C. purpurea* files were used as BUSCO seed species along with the Sordariomyceta database and all five *ab initio* predictions were passed through the --other_gff flag with weights of 1. The following flags were also used in Funannotate “predict”: --repeats2evm, --optimize_augustus, --soft_mask 1000, --min_protlen 50. BUSCO was used to evaluate annotation completeness using the Dikarya and Sordariomyceta databases (odb9) with --prot on default settings.

### Functional annotation

Functional analysis was performed using Funannotate “annotate”. The following analyses were also performed on the three reference *Claviceps* genomes. Secondary metabolite clusters were predicted using antiSMASH v5 (Blin *et al.* 2019) with all features turned on. Functional domain annotations were conducted using eggNOG-mapper v5 (Huerta-Cepas *et al.* 2016, 2019) on default settings and InterProScan v5 (Jones *et al.* 2014) with the --goterms flag. Phobius v1.01 (Käll 2007) was used to assist in prediction of secreted proteins. In addition to these analyses Funannotate also performed domain annotations through an HMMer search against the Pfam-A database and dbCAN CAZYmes database, a BLASTp search against the MEROPS protease database, and secreted protein predictions with SignalP v4.1 (Nielsen 2017).

For downstream analysis, proteins were classified as secreted proteins if they had signal peptides detected by both Phobius and SignalP and did not possess a transmembrane domain as predicted by Phobius and an additional analysis of TMHMM v2.0 (Krogh *et al.* 2001). Effector proteins were identified by using EffectorP v2.0 (Sperschneider *et al.* 2018), with default settings, on the set of secreted proteins for each genome. Transmembrane proteins were identified if both Phobius and TMHMM detected transmembrane domains. Secondary metabolite proteins were identified if they resided within metabolite clusters predicted by antiSMASH. Proteins were classified as having conserved protein domains if they contained any Pfam or IPR domains.

### Gene family identification and classification

OrthoFinder v2.3.3 (Emms & Kelly 2019) was run on default settings using Diamond v0.9.25.126 (Buchfunk *et al.* 2015) to infer groups of orthologous gene clusters (orthogroups) based on protein homology and MCL clustering. To more accurately place closely related genes into clusters an additional 78 fungal genomes (Additional file 1 Table S3) with emphasis on plant associated fungi of the order Hypocreales were added. To standardize, all 78 additional genomes were subject to a protein length cutoff of 50 amino acids and genomes downloaded from http://www.endophyte.uky.edu had alternatively spliced proteins removed. For downstream analysis, orthogroups pertaining to the 53 *Claviceps* genomes were classified as secreted, predicted effectors, transmembrane, metabolite, and conserved domain orthogroups if ≥50% of the *Claviceps* strains present in a given cluster had at least one protein classified as such.

### Phylogeny and genome fluidity

Phylogenetic relationship of all 53 *Claviceps* genomes, with *Fusarium graminearum*, *F. verticillioides*, *Epichloe festucae*, and *E. typhina* as outgroups, was derived from 2,002 single-copy orthologs obtained from our OrthoFinder defined gene clusters (described above). This resulted in a dataset of 114,114 amino acids sequences which were concatenated to create a super-matrix and aligned using MAFFT v7.429 (Katoh & Standely 2013) on default settings. Uninformative sites were removed using Gblocks v0.91 (Castresana 2000) on default settings. Due to the large scale of the alignment maximum likelihood reconstruction was performed using FastTree v2.1.11 (Price *et al.* 2010) using the WAG model of amino acid substitution with the - gamma, -spr 4, -mlacc 2, -slownni, and -slow flag with 1000 bootstraps. MEGA X (Sudhir *et al.* 2018) was used for neighbor-joining (NJ) reconstruction using the JTT model of amino acid substitution with gamma distribution and maximum parsimony (MP) reconstruction using the tree bisection reconstruction (TBR) algorithm with 100 repeated searches. Nodal support for both NJ and MP reconstructions were assessed with 1000 bootstraps. In addition, an alignment and ML reconstruction was performed on each of the 2,002 protein sequences following the procedure as above (MAFFT, Gblocks, FastTree). A density consensus phylogeny was created from all gene trees using the program DensiTree v2.2.5 (Bouckaert & Heled 2014). PhyBin v0.3-1 (Newton & Newton 2013) was used to cluster trees from three datasets (1: *Claviceps* genus without outgroups, 2: sect. *Pusillae* species, 3: sect. *Claviceps* species) together to identify frequencies of concordant topologies using the --complete flag with --editdist=2. To reduce noise, from abundant incomplete lineage sorting in sect. *Claviceps*, we implemented a --minbranchlen=0.015 for our *Claviceps* genus dataset.

Following methodologies established in Kislyuk *et al.* (2011) genomic fluidity, which estimates the dissimilarity between genomes by using ratios of the number of unique gene clusters to the total number of gene clusters in pairs of genomes averaged over randomly chosen genome pairs from within a group on *N* genomes, was used to assess gene cluster dissimilarity within the *Claviceps* genus. For a more detailed description refer to Kislyuk *et al.* (2011). Datasets containing gene clusters from representative members of sect. *Pusillae*, sect. *Claviceps*, *Clavieps* genus, and all *C. purpurea* strains were extracted from our OrthoFinder defined gene clusters. Additional species- and genus-wide gene cluster datasets from the additional 78 fungal genomes were extracted for comparative purposes. All section- and genus-wide datasets contained one representative isolate from each species to reduce phylogenetic bias. Each extracted dataset was used to calculate the genomic fluidity using a custom script (https://github.com/PlantDr430/CSU_scripts/blob/master/pangenome_fluidity.py). The result files for each dataset were then used for figure creation and two-sample two-sided z-test statistics (Kislyuk *et al.* 2011) using a custom script (https://github.com/PlantDr430/CSU_scripts/blob/master/combine_fluidity.py).

### Gene compartmentalization

A custom script (https://github.com/PlantDr430/CSU_scripts/blob/master/genome_speed_hexbins.py) was used to calculate local gene density measured as 5’ and 3’ flanking distances between neighboring genes (intergenic regions). To statistically determine whether specific gene types had longer intergenic flanking regions than all other genes within the genome we randomly sampled 100 each group of genes (specific gene vs. other genes) 1,000 times for both the 5’ and 3’ flanking distances. Mann-Whitney U test was used to test for significance on all 2,000 subsets corrected with Benjamini-Hochberg. Corrected p-values were averaged per flanking side and then together to get a final p-value. Genes that appeared on a contig alone were excluded from analysis. For graphical representation, genes that were located at the start of each contig (5’ end) were plotted along the x-axis, while genes located at the end of each contig (3’ end) were plotted along the y-axis.

### RIP and BLAST analyses

For all 53 genomes a self BLASTp v2.9.0+ search was conducted to identify best hit orthologs within each genome with a cutoff e-value of 10^−5^ and removal of self-hits. This process was automated, using a custom script (https://github.com/PlantDr430/CSU_scripts/blob/master/RIP_blast_analysis.py). We further examined if gene pairs with a pairwise identity of ≥80% were located next to each other and/or separated by five or fewer genes. Fifty-six important *Claviceps* genes (Additional file 2: Table S7) including the *rid-1* homolog (Freitag *et al.* 2002) were used in a BLASTp analysis to identify the number of genes present that passed an e-value cutoff of 10^−5^, 50% coverage, and 35% identity. Genes that appeared as best hits for multiple query genes were only recorded once for their overall best match. In addition, the web-based tool The RIPper (Van Wyk *et al.* 2019) was used on default settings (1 kb windows in 500 bp increments) to scan whole-genomes for presence of RIP and large RIP affected regions (LRARs).

### Statistical programs and plotting

Statistics and figures were generated using Python3 modules SciPy v1.3.1, statsmodel v0.11.0, and Matplotlib v3.1.1. Heatmaps were generated using ComplexHeatmap v2.2.0 in R (Gu 2016).

## Results

### Genome Assembly and Annotation

To provide a comprehensive view of variability across *Claviceps*, we sequenced and annotated 50 genomes (19 *Claviceps* spp.), including *C. citrina* the single species of sect. *Citrinae*, six species belonging to sect. *Pusillae*, and 44 genomes (12 species) belonging to sect. *Claviceps*, of which 23 genomes belong to *C. purpurea* s.s. (Table 1; Additional file 1: Table S1). The assemblies and annotations were of comparable quality to the reference strains (Table 1). A more detailed representation of the assembly and annotation statistics can be seen in Table 1 and Additional file 1: Figure S1, Table S2.

**Table 1:** Assembly and annotations statistics for the three reference *Claviceps* genomes and the 50 *Claviceps* genomes used in this study.

| Organism | Strain | Section | Host of Origin |  | Read Coverage | Genome size (Mb) | Contig (#) | N50 | Genomic GC (%) | TE<sup>§</sup> content (%) | Gene count | BUSCO Completeness |  |
| --- | --- | --- | --- | --- | --- | --- | --- | --- | --- | --- | --- | --- | --- |
|  |  |  | Family / Tribe | Genus / species |  |  |  |  |  |  |  | Dikarya | Sordario-myceta |
| References: |  |  |  |  |  |  |  |  |  |  |  |  |  |
| *C. purpurea* | 20.1 | Claviceps | Triticeae | *Secale cereale* | -- | 32.1 | 1,442<sup>†</sup> | 46,498<sup>†</sup> | 51.6% | 10.9% | 8,703 | 95.30% | 94.70% |
| *C. fusiformis* | PRL1980 | Pusillae | Panicaceae | *Pennisetum typhoideum* | -- | 52.3 | 6,930 | 19,980 | 37.3% | 47.5% | 9,304 | 96.70% | 94.90% |
| *C. paspali* | RRC1481 | Paspalorum | Paspaleae | *Paspalum* sp. | -- | 28.9 | 2,304 | 26,898 | 47.7% | 17.5% | 8,400 | 94.30% | 93.30% |
| This study: |  |  |  |  |  |  |  |  |  |  |  |  |  |
| *C. purpurea* | Clav04 | Claviceps | Bromeae | *Bromus inermis* | 46x | 31.8 | 3,288 | 21,051 | 51.7% | 10.1% | 8,824 | 95.50% | 94.10% |
| *C. purpurea* | Clav26 | Claviceps | Triticeae | *Hordeum vulgare* | 59x | 30.8 | 1,361 | 49,697 | 51.7% | 9.1% | 8,737 | 97.70% | 96.50% |
| *C. purpurea* | Clav46 | Claviceps | Triticeae | *Secale cereale* | 58x | 30.8 | 1,409 | 49,302 | 51.7% | 9.7% | 8,597 | 98.00% | 96.60% |
| *C. purpurea* | Clav55 | Claviceps | Poeae | *Lolium perenne* | 59x | 30.7 | 1,525 | 44,299 | 51.8% | 9.8% | 8,480 | 97.10% | 95.90% |
| *C. purpurea* | LM4 | Claviceps | Triticeae | *Trisetoscale* | 64x | 30.6 | 1,296 | 47,441 | 51.8% | 10.0% | 8,470 | 97.00% | 95.80% |
| *C. purpurea* | LM5 | Claviceps | Triticeae | *Hordeum vulgare* | 67x | 30.5 | 1,258 | 51,505 | 51.8% | 9.0% | 8,508 | 96.90% | 95.50% |
| *C. purpurea* | LM14 | Claviceps | Triticeae | *Hordeum vulgare* | 49x | 30.6 | 1,297 | 49,955 | 51.8% | 10.0% | 8,422 | 97.40% | 95.60% |
| *C. purpurea* | LM28 | Claviceps | Triticeae | *Triticum aestivum* | 49x | 30.6 | 1,343 | 51,635 | 51.7% | 9.6% | 8,713 | 97.30% | 96.10% |
| *C. purpurea* | LM30 | Claviceps | Triticeae | *Secale cereale* | 64x | 30.6 | 1,224 | 51,374 | 51.8% | 9.4% | 8,526 | 97.00% | 95.50% |
| *C. purpurea* | LM33 | Claviceps | Triticeae | *Secale cereale* | 45x | 30.5 | 1,398 | 44,564 | 51.8% | 9.2% | 8,557 | 96.30% | 95.50% |
| *C. purpurea* | LM39 | Claviceps | Triticeae | *Triticum turgidum* subsp. durum | 81x | 30.5 | 1,282 | 48,443 | 51.8% | 10.1% | 8,591 | 97.10% | 96.10% |
| *C. purpurea* | LM46 | Claviceps | Triticeae | *Triticum turgidum* subsp. durum | 79x | 30.6 | 1,291 | 50,932 | 51.8% | 9.6% | 8,455 | 97.00% | 95.80% |
| *C. purpurea* | LM60 | Claviceps | Poeae | *Avena sativa* | 81x | 30.6 | 1,259 | 47,464 | 51.7% | 9.3% | 8,498 | 97.00% | 95.80% |
| *C. purpurea* | LM71 | Claviceps | Poeae | *Alopercurus myosuroides* | 168x | 30.5 | 1,400 | 45,114 | 51.8% | 9.6% | 8,472 | 97.10% | 95.60% |
| *C. purpurea* | LM207 | Claviceps | Triticeae | *Elymus repens* | 53x | 30.5 | 1,352 | 45,388 | 51.8% | 9.2% | 8,475 | 97.00% | 95.70% |
| *C. purpurea* | LM223 | Claviceps | Bromeae | *Bromus riparius* | 74x | 30.8 | 1,297 | 46,577 | 51.7% | 10.5% | 8,438 | 97.00% | 95.70% |
| *C. purpurea* | LM232 | Claviceps | Poeae | *Phalaris canariensis* | 53x | 30.7 | 1,348 | 49,571 | 51.7% | 9.4% | 8,512 | 96.60% | 95.70% |
| *C. purpurea* | LM233 | Claviceps | Poeae | *Phalaris canariensis* | 49x | 30.6 | 1,331 | 50,327 | 51.8% | 9.9% | 8,717 | 96.70% | 95.90% |
| *C. purpurea* | LM461 | Claviceps | Triticeae | *Elymus repens* | 37x | 30.5 | 1,440 | 44,216 | 51.8% | 8.4% | 8,656 | 96.60% | 95.20% |
| *C. purpurea* | LM469 | Claviceps | Triticeae | *Triticum aestivum* | 75x | 30.5 | 1,257 | 48,403 | 51.8% | 10.0% | 8,394 | 97.30% | 96.00% |
| *C. purpurea* | LM470 | Claviceps | Triticeae | *Elymus repens* | 26x | 30.5 | 1,797 | 32,579 | 51.8% | 9.0% | 8,591 | 96.50% | 95.30% |
| *C. purpurea* | LM474 | Claviceps | Triticeae | *Hordeum vulgare* | 64x | 30.6 | 1,354 | 47,245 | 51.8% | 9.4% | 8,500 | 96.80% | 95.70% |
| *C. purpurea* | LM582 | Claviceps | Triticeae | *Secale cereale* | 89x | 30.7 | 1,600 | 39,003 | 51.8% | 9.6% | 8,518 | 97.20% | 95.40% |
| *C. aff. purpurea* | Clav52 | Claviceps | Poeae | *Poa pratensis* | 60x | 29.6 | 1,334 | 48,893 | 51.8% | 8.2% | 8,316 | 96.80% | 96.20% |
| *C. quebecensis*‡ | Clav32 | Claviceps | Triticeae | *Hordeum vulgare* | 64x | 28.7 | 1,068 | 58,118 | 51.6% | 4.5% | 8,232 | 98.00% | 96.60% |
| *C. quebecensis*‡ | Clav50 | Claviceps | Triticeae | *Elymus* sp. | 59x | 28.8 | 1,075 | 66,795 | 51.6% | 6.9% | 8,046 | 97.50% | 96.30% |
| *C. quebecensis*‡ | LM458 | Claviceps | Poeae | *Ammophila* (plant) | 78x | 28.4 | 1,166 | 45,693 | 51.6% | 6.1% | 8,055 | 97.10% | 95.80% |
| *C. occidentalis*‡ | LM77 | Claviceps | Poeae | *Phleum pratense* | 58x | 28.7 | 1,728 | 29,222 | 51.4% | 6.0% | 8,162 | 96.10% | 94.70% |
| *C. occidentalis*‡ | LM78 | Claviceps | Bromeae | *Bromus inermis* | 64x | 28.8 | 1,689 | 29,608 | 51.4% | 6.0% | 8,231 | 95.80% | 94.70% |
| *C. occidentalis*‡ | LM84 | Claviceps | Bromeae | *Bromus inermis* | 164x | 28.9 | 1,404 | 36,685 | 51.4% | 6.0% | 8,221 | 97.00% | 95.40% |
| *C. ripicola*‡ | LM218 | Claviceps | Poeae | *Phalaris arundinacea* | 146x | 31.1 | 1,072 | 60,464 | 51.4% | 10.3% | 8,327 | 96.70% | 95.70% |
| *C. ripicola*‡ | LM219 | Claviceps | Poeae | *Phalaris arundinacea* | 55x | 30.8 | 1,239 | 55,312 | 51.4% | 9.5% | 8,381 | 96.80% | 95.80% |
| *C. ripicola*‡ | LM220 | Claviceps | Poeae | *Phalaris arundinacea* | 91x | 30.9 | 1,223 | 54,100 | 51.4% | 9.3% | 8,449 | 97.10% | 95.90% |
| *C. ripicola*‡ | LM454 | Claviceps | Poeae | *Ammophila breviligulata* | 156x | 31.2 | 1,508 | 40,844 | 51.4% | 8.4% | 8,562 | 97.10% | 96.10% |
| *C. spartinae* | CCC535 | Claviceps | Zoysieae | *Sporobolus anglicus* | 60x | 29.3 | 1,456 | 42,688 | 51.4% | 7.1% | 8,433 | 97.50% | 95.90% |
| *C. arundinis* | LM583 | Claviceps | Molinieae | *Phragmites australis* | 69x | 30.6 | 996 | 70,672 | 51.4% | 9.8% | 8,235 | 96.80% | 95.70% |
| *C. arundinis* | CCC1102 | Claviceps | Molinieae | *Phragmites australis* | 61x | 30.3 | 896 | 91,905 | 51.4% | 8.3% | 8,486 | 97.70% | 96.50% |
| *C. humidiphila* | LM576 | Claviceps | Poeae | *Dactylis* sp. | 77x | 31.2 | 1,236 | 55,717 | 51.5% | 9.9% | 8,440 | 97.00% | 95.90% |
| *C. perihumidiphila*‡ | LM81 | Claviceps | Triticeae | *Elymus albicans* | 140x | 31.2 | 1,003 | 67,487 | 51.5% | 11.0% | 8,291 | 97.10% | 95.90% |
| *C. cyperi* | CCC1219 | Claviceps | Cyperaceae (family) | *Cyperus esculentus* | 56x | 26.6 | 1,921 | 27,113 | 51.7% | 8.9% | 7,673 | 97.70% | 95.40% |
| *C. capensis* | CCC1504 | Claviceps | Ehrharteae | *Ehrharta villosa* | 66x | 27.7 | 1,136 | 59,777 | 51.7% | 6.2% | 8,037 | 97.60% | 95.70% |
| *C. pazoutovae* | CCC1485 | Claviceps | Stipeae | *Stipa dregeana* | 61x | 27.6 | 1,304 | 42,785 | 51.7% | 6.8% | 7,941 | 97.50% | 96.00% |
| *C. monticola* | CCC1483 | Claviceps | Brachypodieae | *Brachypodium* sp. | 58x | 27.8 | 1,144 | 56,619 | 51.6% | 7.0% | 7,977 | 98.10% | 96.50% |
| *C. pusilla* | CCC602 | Pusillae | Andropogoneae | *Bothriochloa insculpta* | 52x | 45.9 | 5,068 | 15,010 | 40.4% | 42.1% | 8,735 | 90.90% | 88.30% |
| *C. lovelessii* | CCC647 | Pusillae | Eragostidinae | *Eragrostis* sp. | 53x | 41.1 | 5,300 | 12,480 | 42.1% | 33.9% | 8,862 | 91.60% | 88.20% |
| *C. digitariae* | CCC659 | Pusillae | Panicaceae | *Digitaria eriantha* | 57x | 33.4 | 1,773 | 32,638 | 44.8% | 20.0% | 8,285 | 95.90% | 94.70% |
| *C. maximensis* | CCC398 | Pusillae | Panicaceae | *Megathyrsus maximus* | 58x | 33.0 | 829 | 81,956 | 44.9% | 19.8% | 7,943 | 98.30% | 96.50% |
| *C. sorghi* | CCC632 | Pusillae | Andropogoneae | *Sorghum bicolor* | 60x | 35.6 | 3,660 | 16,225 | 44.4% | 30.4% | 8,208 | 89.90% | 87.10% |
| *C. africana* | CCC489 | Pusillae | Andropogoneae | *Sorghum bicolor* | 56x | 37.7 | 1,781 | 37,639 | 42.5% | 34.0% | 8,119 | 95.00% | 91.50% |
| *C. citrina* | CCC265 | Citrinae | Cynodonteae | *Distichlis spicata* | 64x | 43.5 | 4,772 | 16,294 | 41.5% | 51.7% | 7,821 | 92.20% | 88.20% |

† The reference strain *C. purpurea* 20.1 was additionally assembled into 191 scaffolds with a scaffold N50 of 433,221.
‡ Newly identified species Liu *et al.* (Submitted)
§ Transposable element (TE) content represented as percent of the genome masked by TEs

Overall, species of sect. *Claviceps* had better assemblies and annotations than species of other sections regarding contig numbers, N50’s, and BUSCO completeness scores (Table 1). Nearly all species of sect. *Claviceps* showed higher BUSCO scores than the references, while species of sects. *Pusillae* and *Citrinae* generally showed lower scores, likely due to their higher TE content (average 34.9% ± 11.0%; Table 1). Exceptions to the low BUSCO scores were *C. digitariae* and *C. maximensis* (sect. *Pusillae*), which had lower TE content, 20.0% and 19.8%, respectively, than the rest of the species in sect. *Pusillae* (Table 1).

### Phylogenomics and genome fluidity

Orthologous gene clusters (orthogroups), which contain orthologs and paralogs, were inferred from protein homology and MCL clustering using OrthoFinder. Across the 53 *Claviceps* isolates and outgroups species *Fusarium graminearum*, *F. verticillioides*, *Epichloe festucae*, and *E. typhina*. We identified 2,002 single-copy orthologs. We utilized a super-matrix approach to infer a maximum likelihood (ML) species tree, based on these protein sequences. Results showed statistical support for four sections of *Claviceps* with a near concordant topology to the Bayesian five-gene phylogeny in Píchová *et al.* (2018). This topology was supported by neighbor-joining and maximum parsimony super-matrix analyses (Additional file 1: Figure S2, S3). Notable exceptions were the placement of *C. paspali* (sect. *Paspalorum*) which grouped closer to *C. citrina* (sect. *Citrinae*) instead of sect. *Claviceps,* and *C. pusilla* which grouped closer to *C. fusiformis* instead of *C. maximensis* (Fig. 1). We also found that sect. *Claviceps* diverged from a common ancestor with sect. *Pusillae* as opposed to sect. *Paspalorum*. Our results provide support for the deeply divergent lineages of sects. *Pusillae*, *Paspalorum*, and *Citrinae* with a long divergent branch resulting in sect. *Claviceps* (Fig. 1).

**Figure 1:**
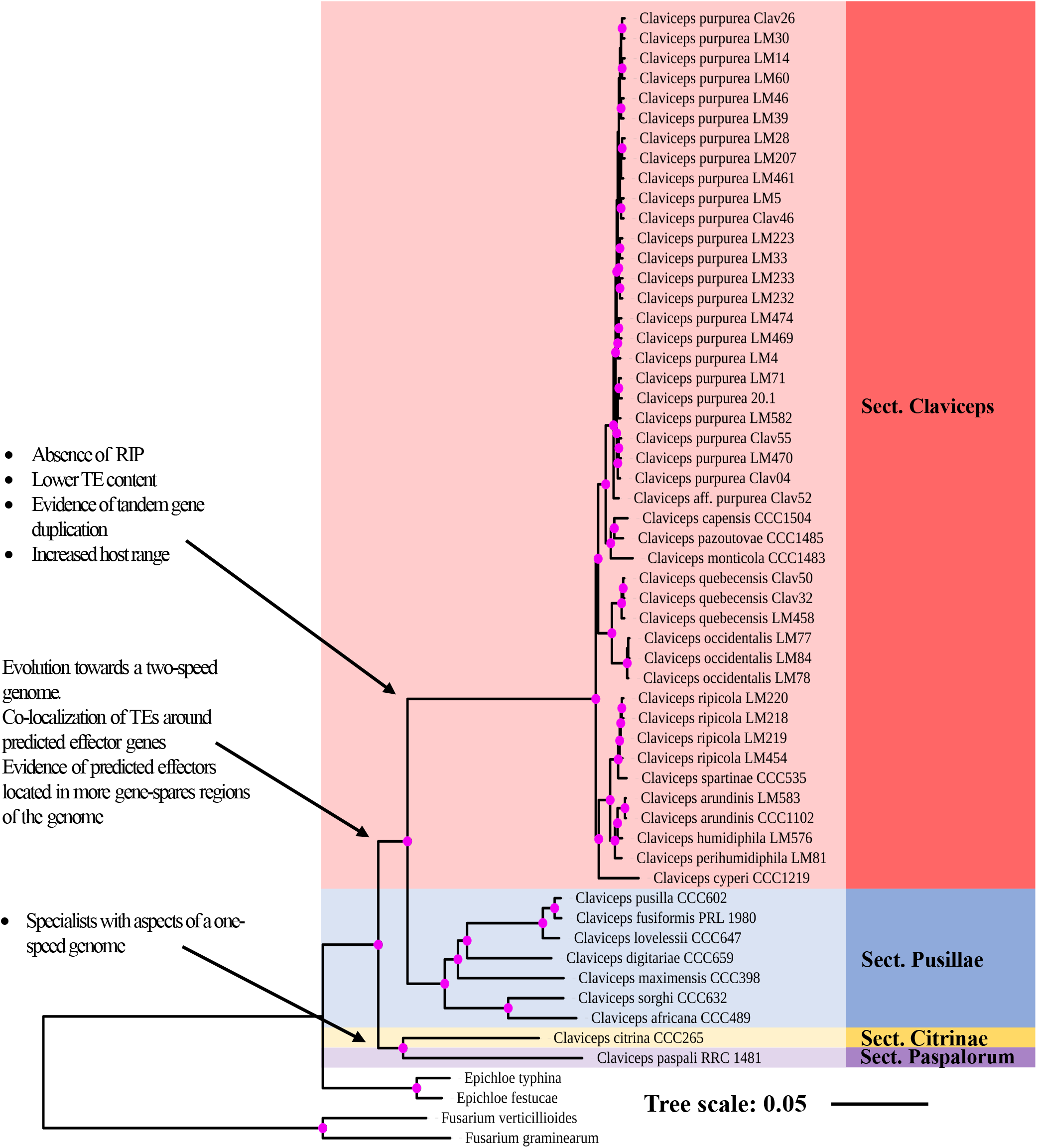
Maximum likelihood phylogenetic reconstruction of the *Claviceps* genus using amino acid sequences of 2,002 single copy orthologs with 1000 bootstrap replicates. Pink dots at branches represent bootstrap values ≥95. Arrows and descriptions indicate potential changes in genomic architecture between *Claviceps* sections identified in this study.

Each of the 2,002 single-copy orthologs were also independently aligned and analyzed in the same manner as our super-matrix phylogeny from representative isolates of each species. A density consensus tree of all 2,002 topologies was concordant with our super matrix analysis but reveals evidence of incongruencies, particularly within sect. *Claviceps* (Additional File 1 Fig. S4) which could be caused by biological, analytical, and sampling factors (Steenwyk *et al.* 2019). While grouping of species generally held true to Fig. 1, variation was more related to the order of branches, with *C. cyperi*, *C. arundinis*, *C. humidiphila*, and *C. perihumidiphila* showing the most variability. These results indicate the presence of some incongruencies within sect. *Claviceps*, sect. *Pusillae*, and across the genus (Additional File 1 Fig. S5-7) but a consensus supporting our ML species tree (Fig. 1, Additional File 1 Fig. S4). There are several potential causes of these incongruencies which are currently the focal point of an ongoing study.

To further elucidate trends of divergence within the genus we examined genomic fluidity (Kislyuk *et al.* 2011) using all 82,267 orthogroups from our previous OrthoFinder analysis. Genomic fluidity estimates the dissimilarity between genomes by using ratios of the number of unique orthogroups to the total number of orthogroups in pairs of genomes averaged over randomly chosen genome pairs from within a group on *N* genomes. For example, a fluidity value of 0.05 indicates that randomly chosen pairs of genomes in a group will on average have 5% unique orthogroups and share 95% of their orthogroups (Kislyuk *et al.* 2011). Section *Claviceps*, which is composed of 12 different species, showed a relatively small genomic fluidity (0.0619 ± 0.0019) with limited variation, indicating pairwise orthogroup dissimilarity between randomly sampled genomes was quite low. The amount of variation between 12 different *Claviceps* species was similar to the variation between 24 *C. purpurea* s.s. isolates, however, the fluidities were significantly different (Additional file 1: Table S4; *P* < 0.0001). In comparison, the fluidity of sect. *Pusillae* (0.126 ± 0.014; *P* < 0.0001) was two times greater than the fluidity of sect. *Claviceps* and exhibited greater variation, indicating greater dissimilarities in orthogroups between randomly sampled species of sect. *Pusillae*.

Overall, our ML phylogeny (Fig. 1) and genome fluidity analysis (Fig. 2) indicate a large evolutionary divergence separating sect. *Claviceps*. Our subsequent analyses of the genomic architecture of all *Claviceps* species examine factors that could be associated with the evolutionary divergence of sect. *Claviceps* and those driving cryptic speciation.

**Figure 2:**
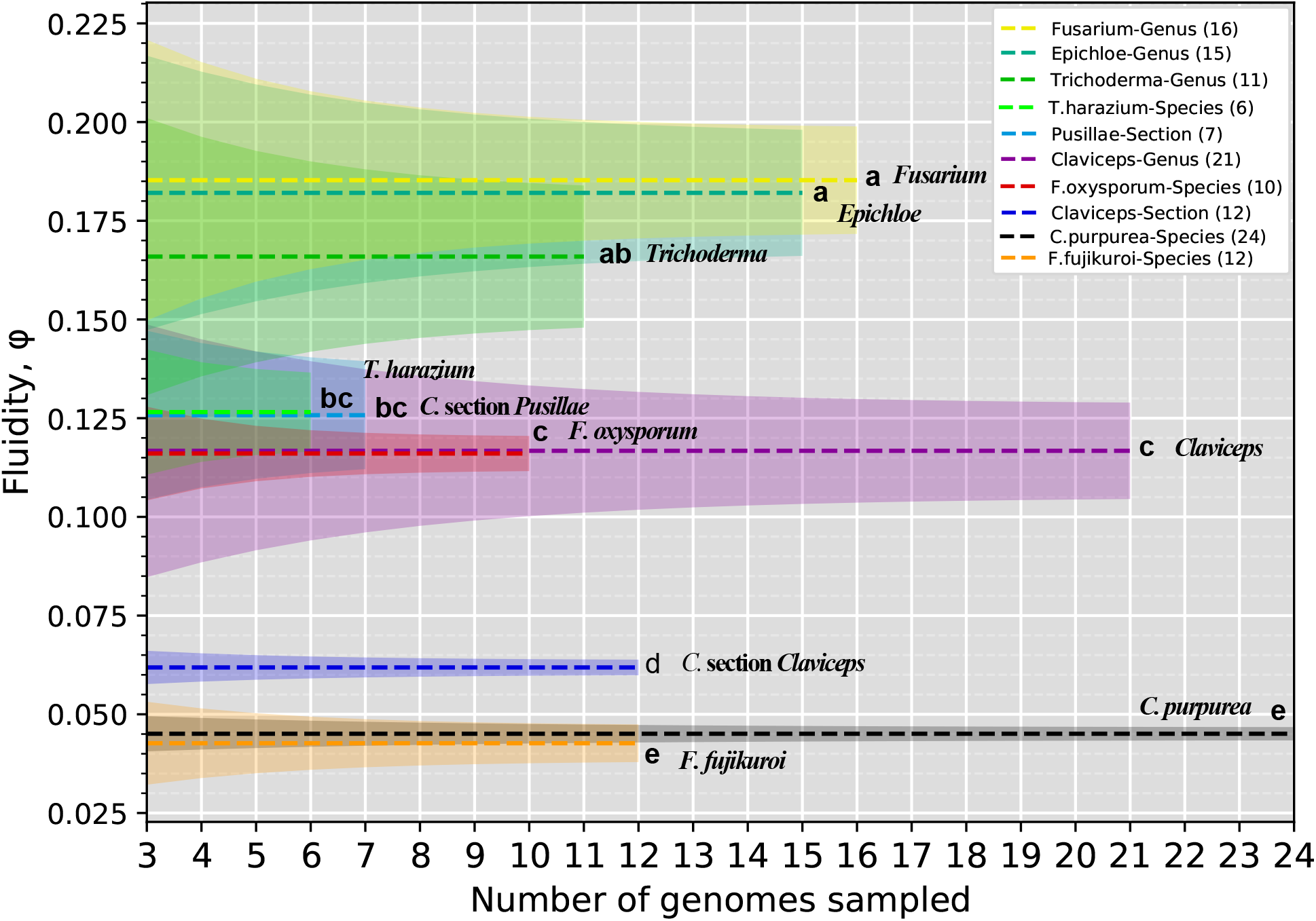
Genomic fluidity (dashed lines) for specified groups within the order Hypocreales. Species level groups contain multiple isolates of a given species, while section and genus level groups contain one strain from representative species to remove phylogenetic bias. Shaded regions represent standard error and were determined from total variance, containing both the variance due to the limited number of samples genomes and the variance due to subsampling within the sample of genomes. Letters correspond to significant difference between fluidities determined through a two-sided two-sample z-test (*P* < 0.05; Additional file 1: Table S4). Legend is in descending order based on fluidity, and names are additionally appended to mean lines for clarity.

### Transposable element divergences and locations

Transposable element (TE) divergence landscapes revealed an overrepresentation of LTR elements in sects. *Pusillae*, *Citrinae*, and *Paspalorum*. All three sections showed a similar large peak of LTRs with divergences between 5-10% (Fig. 3; Additional file 1: Figure S8), indicating a relatively recent expansion of TEs. The landscapes of sects. *Pusillae*, *Citrinae*, and *Paspalorum* are in striking contrast to species of sect. *Claviceps* which showed more similar abundances of LTR, DNA, LINE, SINE, and RC (helitron) elements. Species of sect. *Claviceps* showed broader peaks of divergence between 5-30% but also showed an abundance of TEs with ~ 0% divergence suggesting very recent TE expansion (Fig. 3; Additional file 1: Figure S8). The TE landscape of *C. cyperi* showed a more striking peak of divergence between 5-10% that more closely resembled the TE divergences of sects. *Pusillae*, *Paspalorum*, and *Citrinae*. However, the content of the TE peak in *C. cyperi* largely contained DNA, LINE, and unclassified TEs as opposed to LTR’s (Additional file 1: Figure S8).

**Figure 3:**
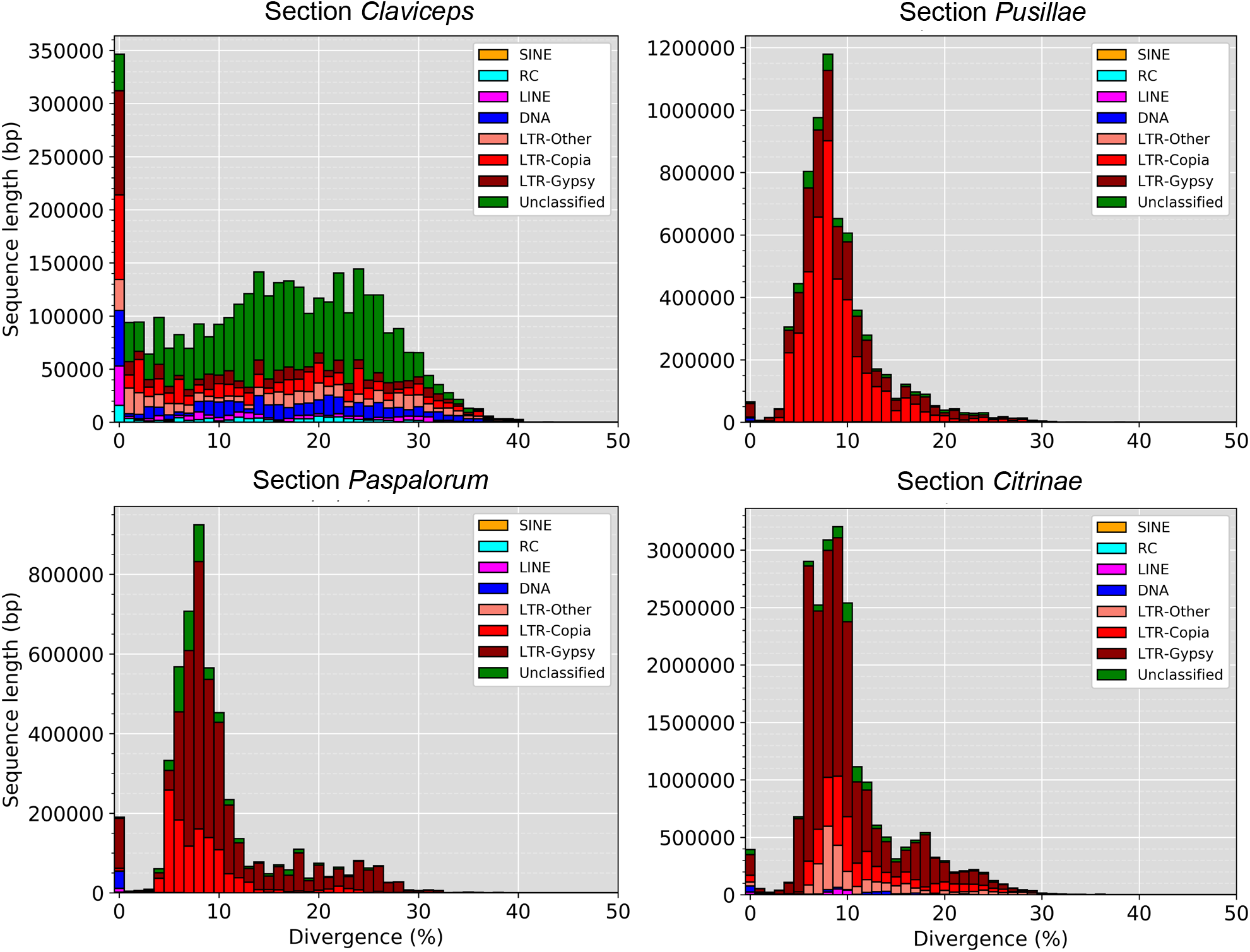
Transposable element (TE) fragment divergence landscapes for representative species of each *Claviceps* section; *C. purpurea* 20.1 (sect. *Claviceps*), *C. maximensis* CCC398 (sect. *Pusillae*), *C. paspali* RRC1481 (sect. *Paspalorum*), and *C. citrina* (sect. *Citrinae*). Stacked bar graphs show the non-normalized sequence length occupied in each genome (y-axis) for each TE type based on their percent divergence (x-axis) from their corresponding consensus sequence. Landscape for all remaining isolates can be seen in Additional file 1: Figure S8.

To identify where genes were located in relation to TEs, we calculated the average distance (kbp) of each gene to the closest TE fragment. This analysis was performed for predicted effectors, secreted (non-effector) genes, secondary metabolite (non-secreted) genes, and all other genes. Secreted genes and predicted effectors of sects. *Claviceps* and *Pusillae* species were found to be significantly closer to TEs compared to other genes within each respective section (Fig. 4; *P* < 0.05), suggesting that these genes could be located in more repeat-rich regions of the genome. It should be noted that we did observe a significant difference (*P* < 0.001, Welch’s test) in TE content between sect. *Pusillae* (32.5% ± 9.59%) and sect. *Claviceps* (8.79.28% ± 1.52%). In both sects. *Claviceps* and *Pusillae* secondary metabolite genes were located farther away from TEs (Fig. 4; *P* < 0.05), i.e. repeat-poor regions of the genome. These trends hold true for individual isolates, with a notable exception of *C. pusilla* (sect. *Pusillae*) showing no significant differences in the proximity of TEs to specific gene types (Additional file 1: Figure S9; *P* > 0.05). Variation existed in whether particular isolates had significant differences between all other genes compared to secreted genes and secondary metabolite genes, but all species in sects. *Claviceps* and *Pusillae* (aside from *C. pusilla*) had predicted effector genes located significantly closer to TEs (Additional file 1: Figure S9; *P* < 0.05). No significant differences in the proximity of TEs to specific gene types were observed in sects. *Citrinae* and *Paspalorum* (Fig. 4; *P* > 0.05), suggesting that TE’s are more randomly distributed throughout these genomes.

**Figure 4:**
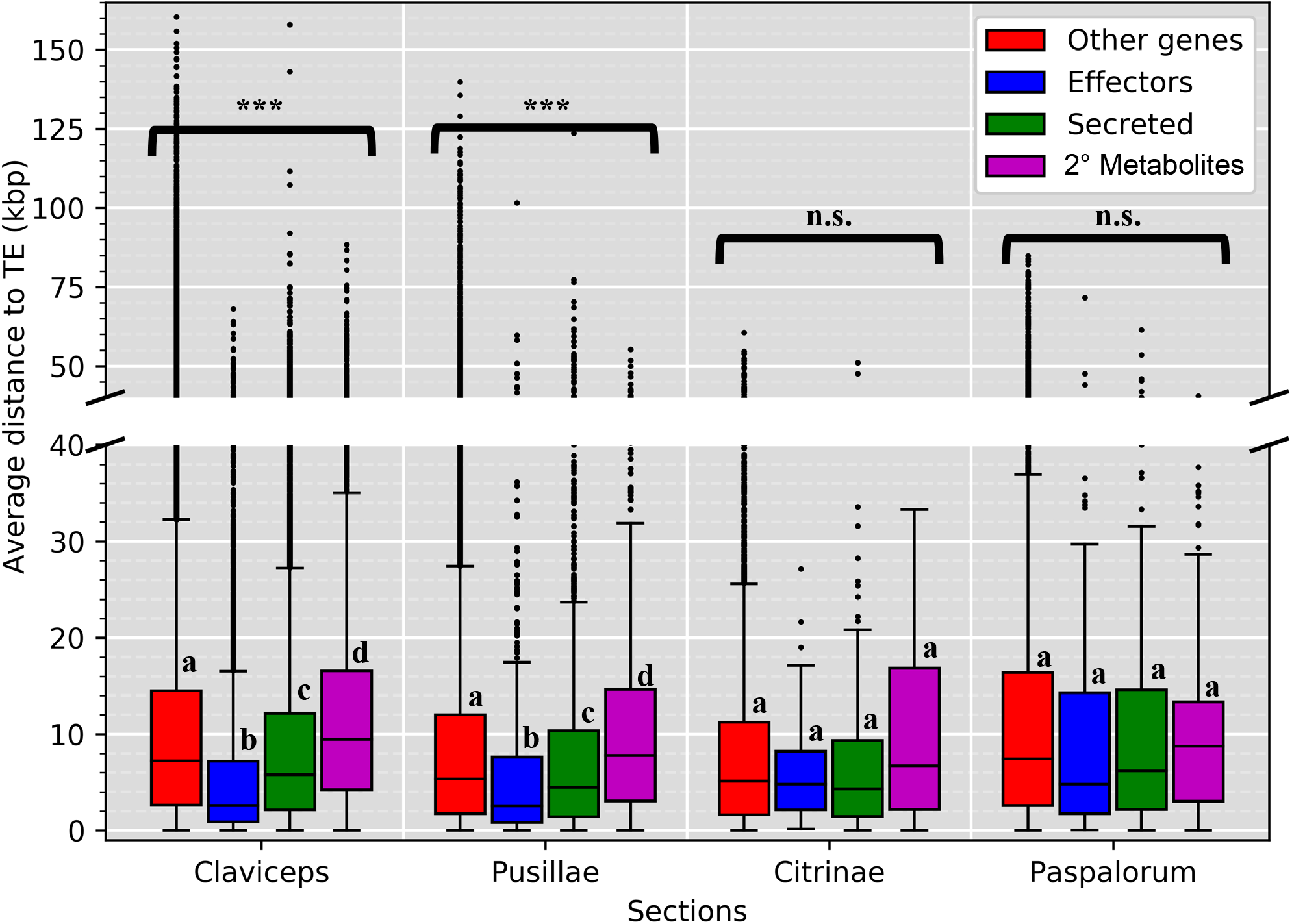
Boxplot distributions of predicted effectors, secreted (non-effectors), secondary metabolite (non-secreted) genes and other genes (i.e. genes that are not effectors, secreted, or secondary (2°) metabolite genes) in *Claviceps* sections showing the mean distance (kbp) of each gene to the closest transposable element fragment (5’ and 3’ flanking distances were averaged together). Kruskal Wallis (*P*-value; * < 0.05, ** < 0.01, *** < 0.001, n.s. = not significant). Pairwise comparison was performed with Mann-Whitney U-test with Benjamini-Hochberg multi-test correction. Letters correspond to significant differences between gene categories within sections (*P* < 0.05). Plots for all individual isolates can been seen in Additional file 1: Figure S9.

### Genome compartmentalization

To further examine genome architecture, we analyzed local gene density measured as flanking distances between neighboring genes (intergenic regions) to examine evidence of genome compartmentalization (i.e. clustering of genes with differences in intergenic lengths) within each genome. Results showed that all 53 *Claviceps* strains exhibited a one-compartment genome (lack of large-scale compartmentalization). Although, there was a tendency for more genes with larger intergenic regions in sects. *Claviceps* and *Pusillae* compared to sects. *Citrinae* and *Paspalorum* (Fig. 5; Additional file 1: Figure S10).

**Figure 5:**
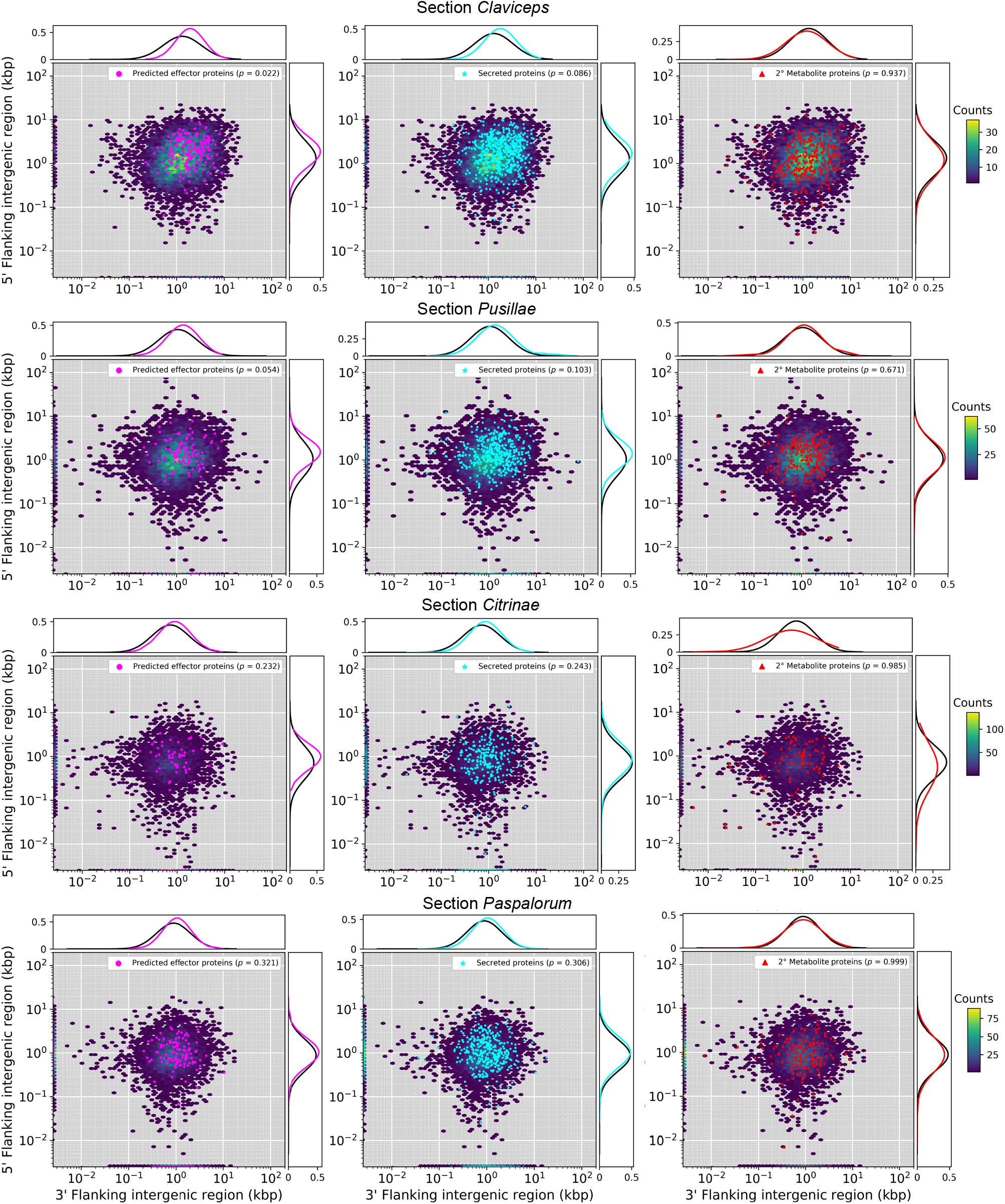
Gene density as a function of flanking 5’ and 3’ intergenic region size (y- and x-axis) of representative isolates of each of the four sections within the *Claviceps* genus; *C, purpurea* 20.1 (sect. *Claviceps*), *C. maximensis* CCC398 (sect. *Pusillae*), *C. paspali* RRC1481 (sect. *Paspalorum*), and *C. citrina* (sect. *Citrinae*). Colored hexbins indicate the intergenic lengths of all genes with color-code indicating the frequency distribution (gene count) according to the legend on the right. Overlaid markers indicate specific gene types corresponding to legends in the top right within each plot. Line graphs (top and right of each plot) depict the frequency distributions of specific gene types (corresponding legend color) and all other genes not of the specific type (black). For visualization purposes the first genes of contigs (5’ end) are plotted along the x-axis and the last gene of each contig (3’ end) are plotted along the y-axis. For information on statistical test see Methods and for plots of all remaining isolates see Additional file 1: Figure S10.

To further clarify evolutionary tendencies, we evaluated whether gene types showed a difference in their flanking intergenic lengths compared to other genes within their genomes. Results showed that predicted effector genes in sect. *Claviceps* had significantly larger intergenic flanking regions compared to other genes, indicating they may reside in more gene-sparse regions of the genome (*P* < 0.05, Fig. 5, Additional file 1: Figure S10). Only *C. digitariae* and *C. lovelessi* (Additional file 1: Figure S10; *P* < 0.01, *P* = 0.024, respectively) of sect. *Pusillae* had predicted effector genes with significantly l arger intergenic regions than other genes, although, *C. fusiformis* and *C. pusilla* were near significant (Fig. 5, Additional file 1: Figure S10; *P* = 0.054, *P* = 0.056, respectively). Flanking intergenic lengths of secreted genes also showed larger intergenic lengths and were often significantly larger than other genes in sect. *Claviceps* (Fig. 5; Additional file 1: Figure S10). In contrast, secondary metabolite genes exhibited a widespread distribution of intergenic lengths that were not significantly different than other genes in all 53 *Claviceps* strains (*P* > 0.05, Fig. 5; Additional file 1: Figure S10).

### RIP analysis

To test for effects of RIP, we assessed the bi-directional similarity of genes against the second closest BLASTp match within each isolate’s own genome (Galagan *et al.* 2003; Urguhart *et al.* 2018), supported by a BLASTp analysis against the *rid-1* RIP gene of *Neurospora crassa*, and calculations of RIP indexes in 1 kb windows (500 bp increments) using The RIPper (Van Wyk *et al.* 2019). Results showed that sects. *Pusillae*, *Citrinae*, and *Paspalorum* had homologs of *rid-1*, fewer genes with close identity (≥ 80%), on average 27.4% ± 11.4% of their genomes affected by RIP, a mean RIP composite index of −0.03 ± 0.21, and 325 ± 138 large RIP affected regions (LRAR) covering 3,984 kb ± 2,144 kb of their genomes, indicating past or current activity of RIP (Fig. 6; Additional file 1: Table S5, S6; Additional file 2: Table S7). This is further supported by an average GC content of 42.84% ± 3.03% (Table 1) in sects. *Pusillae*, *Citrinae*, and *Paspalorum*, which is on average 8.81% lower than in sect. *Claviceps* which shows an absence of RIP (reported below). The presence of RIP in sects. *Pusillae*, *Citrinae*, and *Paspalorum* was unexpected given the abundance of TEs within genomes of these sections (Table 1; Fig. 3; Additional file 1: Figure S8) as RIP should be working to silence and inactivate these TEs. While we did not directly test the activity of TEs within our genomes, due to lack of RNAseq data, the peaks of low TE nucleotide divergence (<10%) in sects. *Pusillae*, *Citrinae*, and *Paspalorum* (Fig. 3, Additional file 1 Figure S8) suggest recent activity of TEs (Frantzeskakis *et al.* 2018).

**Figure 6:**
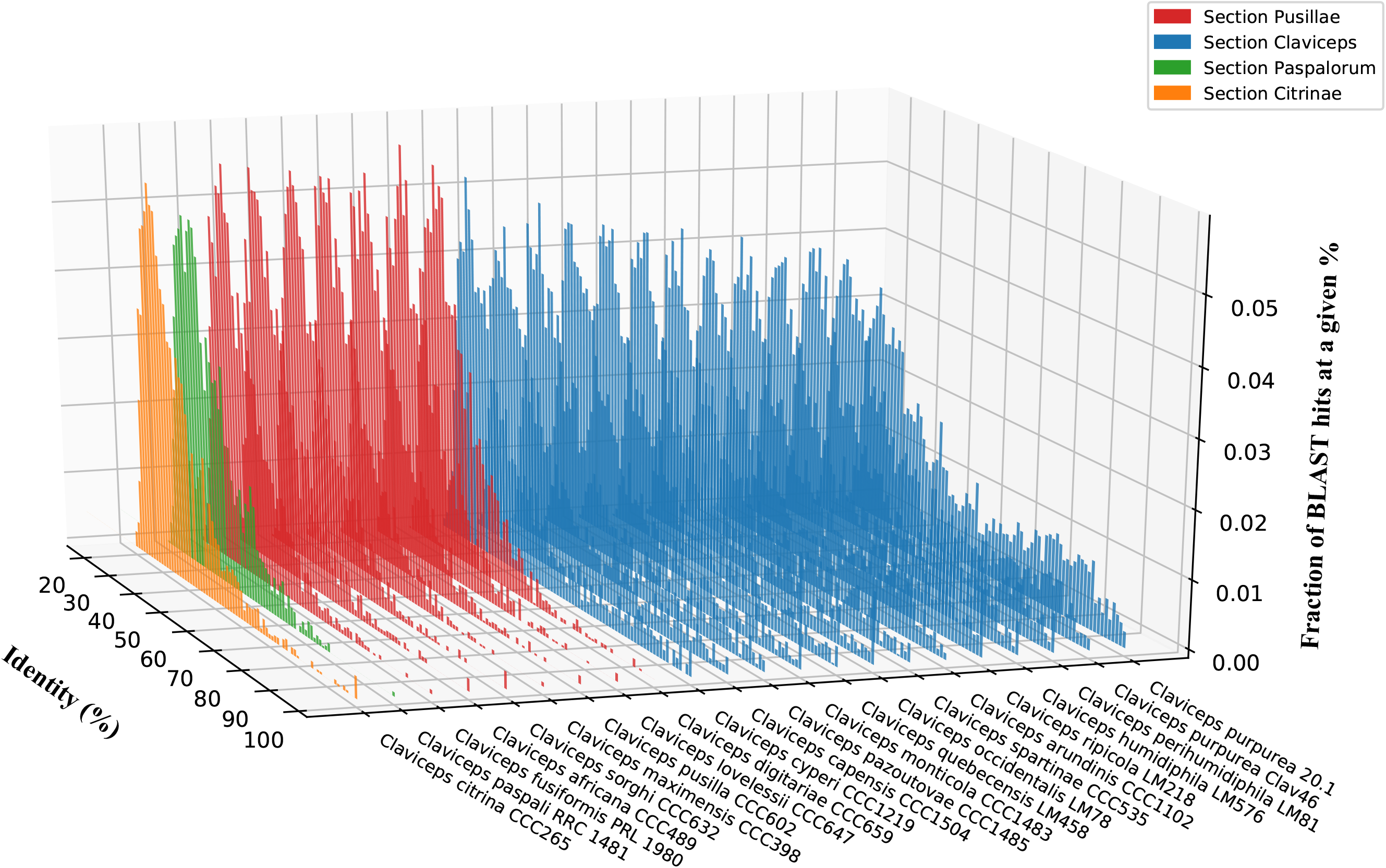
Representative isolates of each *Claviceps* species showing the fraction of BLAST hits at a given % identity (y-axis) within each isolate (z-axis) at a given percent identity (x-axis) from the second closet BLASTp match of proteins within each isolate’s own genome. Two *C. purpruea* s.s. isolates are shown to compare a newly sequenced genome versus the reference.

In comparison, species in sect. *Claviceps* lack *rid-1* homologs, showed larger amounts of gene similarity, and a general lack of evidence of RIP with only 0.13% ± 0.03% of their genomes putatively affected by RIP, and a mean RIP composite index of −0.59 ± 0.01 suggesting that RIP is inactive (Fig. 6; Additional file 1: Table S5, S6; Additional file 2: Table S7). Gene pairs sharing a ≥80% identity to each other were often located near each other. On average 27.02% ± 5.91% of the pairs were separated by five or fewer genes, and 15.95% ± 3.50% of the pairs were located next to each other, indicating signs of tandem gene duplication within the section (Additional file 1: Table S5). *Claviceps cyperi* showed the smallest proportions of highly similar tandem genes (7.77% and 5.7%) compared to other species within sect. *Claviceps*. Additional variations in the proportions of highly similar tandem genes between other species of sect. *Claviceps* were not evident as these proportions appeared to vary more between isolate than species (Additional file 1: Table S5).

### Gene cluster expansion

The proteome of *Claviceps* genomes were used to infer orthologous gene clusters (orthogroups) through protein homology and MCL clustering using OrthoFinder. Our results revealed evidence of orthogroup expansion within sect. *Claviceps* as species contained more genes per orthogroup than species of the other three sections (Additional file 1: Figure S11). To identify the types of gene clusters that were showing putative expansion we filtered our clusters by two criteria; 1) at least one isolates had two or more genes in the orthogroup, 2) there was a significant difference in the mean number of genes per orthogroup between all 44 isolates in sect. *Claviceps* and the 9 isolates from sects. *Pusillae*, *Citrinae*, and *Paspalorum* (α ≤ 0.01, Welch’s test).

Overall, we identified 863 (4.7%) orthogroups showing putative expansion. We observed extensive expansion (orthogroups with observations of ≥ 10 genes per isolate) present in many unclassified, predicted effectors, secreted (non-effector) orthogroups, and orthogroups encoding genes with conserved domains (Fig. 7; Additional file 1: Figure S12, S13). Transmembrane orthogroups also showed evidence of expansion with several isolates having 5-10 genes. orthogroups with secondary metabolite genes showed the lowest amount of expansion (Fig. 8). Overall, sect. *Claviceps* showed expansion in a greater number of orthogroups than sect. *Pusillae*, *Citrinae*, and *Paspalorum* in all categories except transmembranes (Additional file 1: Figure S14). Orthogroups with an average ≥ 5 genes per isolate, within sect. *Claviceps*, contained a variety of functional proteins, with generally more proteins encoding protein/serine/tyrosine kinase domains (Additional file 2: Table S8). Additional details can be obtained from Additional file 2: Table S9 (ordered orthogroups corresponding to heatmaps; Fig. 7 and Additional file 1: Figure S12, S13) and Additional file 3: Table S10-1,2 (orthogroups identification and functional annotation of all proteins).

**Figure 7:**
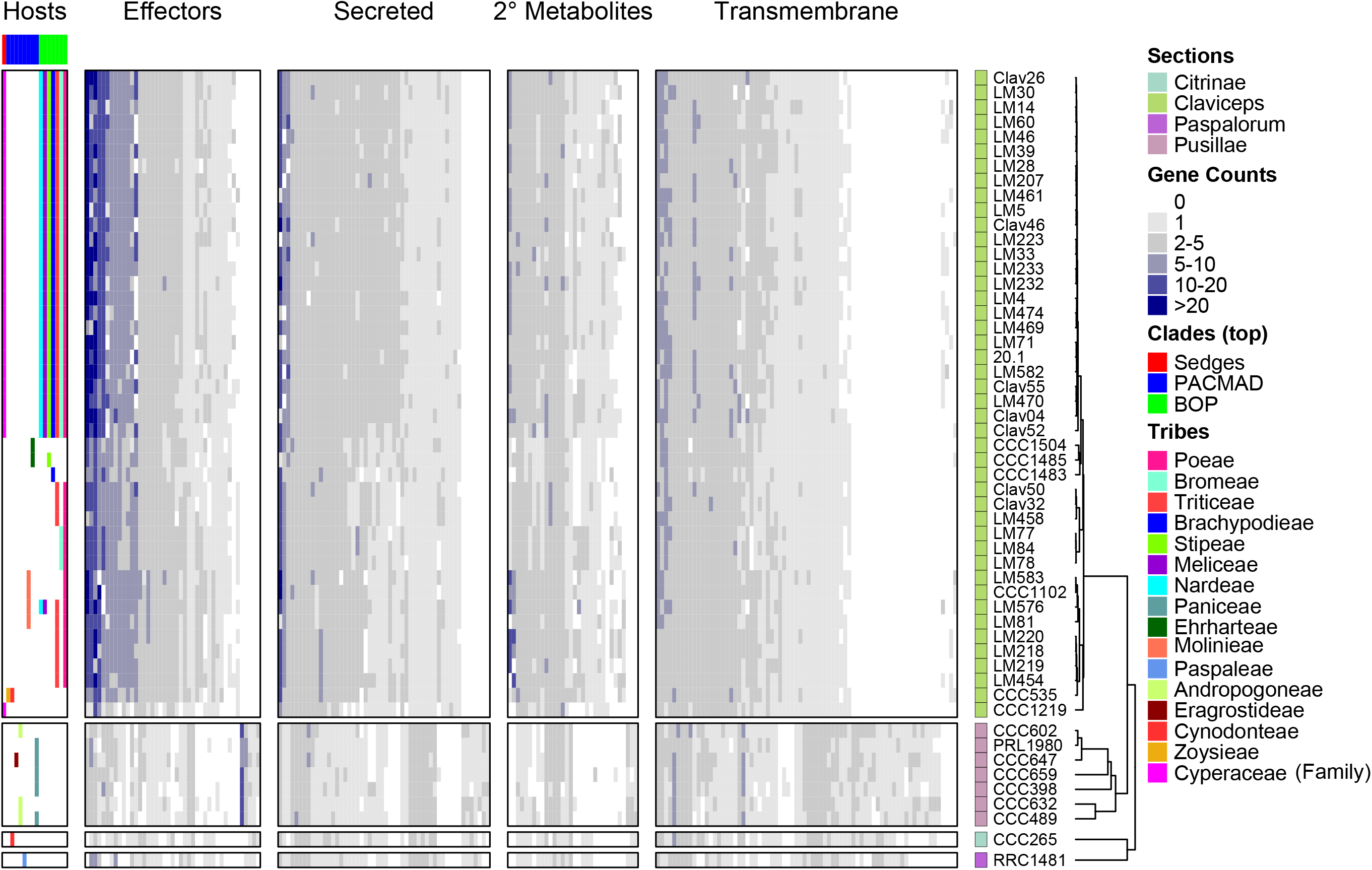
Heatmap of gene counts in orthogroups for all 53 *Claviceps* strains ordered based on ML tree in Fig. 1 and separated by sections. Orthogroups are separated based on their classification and are only represented once (i.e. secondary (2°) metabolite orthogroups shown are those that are not already classified into the effector or secreted orthogroups) and are ordered based on hierarchical clustering, see Additional file 2: Table S9 for list of orthogroups corresponding to the order shown in the heatmaps. The host spectrum (right) is generalized across species, as no literature has determined the existence of race specific isolates within species, is shown on the left side of the figure determined from literature review of field collected samples (Supplementary Material in Píchová *et al.* 2018) and previous inoculation tests Campbell (1957) and Liu *et al.* (*Submitted*). For heatmap of conserved domains see Additional file 1: Figure S12 and for unclassified gene families see Additional file 1: Figure S13.

Within sect. *Claviceps* patterns of gene counts per orthogroup appeared to break down and contain variations in the number of genes per orthogroups with some presence/absences occurring between isolates and species. Notably, *C. cyperi* (CCC1219) showed the lowest amount of expansion, across all taxa, in comparison to other species of sect. *Claviceps*. In addition, *C. spartinae* (CCC535)*, C. capensis* (CCC1504), *C. monticola* (CCC1483), *C. pazoutovae* (CCC1485), *C. occidentalis* (LM77, 78, 84), and *C. quebecensis* (LM458, Clav32, 50) also showed lower expansion (Fig. 7, Additional file 1: Figure S12, S13). However, no patterns were observed linking the variation in expansions with the literature determined host range of different species within sect. *Claviceps*.

## Discussion

Our comparative study of 50 newly annotated genomes from four sections of *Claviceps* has provided us with an enhanced understanding of evolution in the genus through knowledge of factors contributing to its diversification. Our results have revealed that despite having nearly identical life-strategies, these closely related species have substantially altered genomic architecture and plasticity, which may drive genome adaptation. One key difference we observe is a shift from characteristic aspects of one-speed genomes (i.e. less adaptable) in narrow host-range *Claviceps* species (sects. *Citrinae* and *Paspalorum*) towards aspects of two-speed genomes (i.e. more adaptable) in broader host-range lineages of sects. *Pusillae* and *Claviceps* (Fig. 1) (Dong *et al.* 2015; Frantzeskakis *et al.* 2019).

The basal species of the genus, *C. citrina* (sect. *Citrinae*) and *C. paspali* (sect. *Paspalorum*), are characterized by a proliferation of TEs, particularly LTRs, which do not appear to be co-localized around particular gene types (Fig. 4). Coupled with a lack of large-scale genome compartmentalization (Fig. 5), these two species can be considered to fit the concept of a one-speed genome which are often considered to be less adaptable and potentially more prone to being purged from the biota (Dong *et al.* 2015; Frantzeskakis *et al.* 2019). This could help explain the paucity of section lineages and restricted host range to one grass tribe, as similar patterns of large genome size, abundant TE content, and equal distribution of TEs has been observed in the specialized barley pathogen *Blumeria graminis* f.sp. *hordei* (Frantzeskakis *et al.* 2018). Although, rapid adaptive evolution within *B. graminis* f.sp. *hordei*, has been suggested to occur through copy-number variation and/or heterozygosity of effector loci (Dong *et al.* 2015; Frantzeskakis *et al.* 2018, 2019). Our results show a lack of gene duplication occurring in sects. *Citrinae* and *Paspalorum*, likely due to the presence of RIP. However, even with the presence of RIP there was a high LTR content in these species (Fig. 3). This suggests that these LTR elements have found a way to avoid RIP or indicate that these species harbor a less active version of RIP as is found in several fungal species (Kachroo *et al.* 1994; Nakayashiki *et al.* 1999; Graïa *et al.* 2001; Ikeda *et al.* 2002; Chalyet *et al.* 2003; Kito *et al.* 2003). Nonetheless, due to the high abundance of TEs (Fig. 4) and presence of RIP (Fig. 6; Additional file 1: Table S5, S6), we hypothesize that aspects of RIP “leakage” could be a likely mechanism for evolution in *C. citrina* and *C. paspali* (and similarly sect. *Pusillae*), as has been shown to occur in other fungi (Fudal *et al.* 2009; Van de Wouw *et al.* 2010; Hane *et al.* 2015). It should be noted that since the estimated divergence of sect. *Citrinae* 60.5 Mya (Píchová *et al.* 2018) it has remained monotypic. It was only recently that unknown lineages of sect. *Paspalorum* were identified (Oberti *et al.* 2020), although, these lineages were found on the same genera of host as *C. paspali* (*Paspalum* spp.) supporting our hypothesis that species within sect. *Paspalorum* have restricted host ranges. These recent findings further suggest that lack of additional lineages within these sections could be due to limited records of *Claviceps* species in South America, where the genus is thought to have originated (Píchová *et al.* 2018). Further research into South American populations of *Claviceps* will provide significant insight into the evolution of these two sections.

Members of sect. *Pusillae* also exhibited a proliferation of TEs, however, as this section diverged from sects. *Citrinae* and *Paspalorum* the genomic architecture evolved such that TEs co-localized around predicted effector genes (Fig. 4). This proximity of TEs to effectors persisted in sect. *Pusillae* species (except *C. pusilla*; Additional file 1: Figure S9) and sect. *Claviceps* species potentially resulting in the large intergenic regions flanking predicted effector genes (Fig. 5, Additional file 1: Figure S10). Together, these genomic alterations indicate aspects of a two-speed genome (Dong *et al.* 2015; Möller & Stukenbrock 2017). We hypothesize that these observed genomic changes influenced the divergence and adaptability of sects. *Pusillae* and *Claviceps* (Fig. 1) (Raffaele & Kamoun 2012; Stukenbrock 2013; Möller & Stukenbrock 2017).

Furthermore, our analyses suggest that the divergence of sect. *Claviceps*, from sect. *Pusillae*, is associated with a loss of RIP (Fig. 1, 6; Additional file 1: Table S6). In the absence of RIP, the gene-sparse regions rich in TEs and effectors could be hot spots for duplication, deletion, and recombination (Galagan *et al.* 2003, 2004; Raffaele & Kamoun 2012; Dong *et al.* 2015; Faino *et al.* 2016; Möller & Stukenbrock 2017; Frantzeskakis *et al.* 2018, 2019). This would explain the observations of tandem gene duplication within the section (Fig. 6, 7; Additional file 1: Table S5, Figure S11-S14), which may facilitate rapid speciation, as has been postulated in several smut fungi (Kämper *et al.* 2006; Schirawski *et al.* 2010; Dutheil *et al.* 2016). In fact, *C. cyperi*, a basal species of sect. *Claviceps* and thought to be ancestral from ancestral state reconstructions of host range (Píchová *et al.* 2018), showed the least amount of gene cluster expansion and tandem duplication (Fig. 7; Additional file 1: Table S5, Figure S12, S13). Potentially indicating that gene duplication is contributing to the divergence of new species, as other species in sect. *Claviceps* have increased genome size, gene count, and number of closely related gene pairs (≥80% identity) (Table 1; Additional file 1 Table S5). Within sect. *Claviceps* gene duplication is likely facilitated by recombination events during annual sexual reproduction (Esser & Tudzynski 1978). Future studies on recombination will be critical to our understanding of the mechanisms driving gene duplication and elucidating factors associated with the observations of potential incomplete lineage sorting (Pease & Hahn 2013) within the section.

Substantially altered genomic architecture and plasticity between *Claviceps* sections was observed in this study, yet it is unclear whether the evolution of these genomes were caused by contact with new hosts and different climates as ancestral lineages migrated out of South America (Píchová *et al.* 2018), or if the evolution towards a two-speed genome provided an advantage in adapting to new hosts or environments. Further research is needed to clarify this point. As sects. *Pusillae* and *Claviceps* have larger host ranges (5 tribes and 13 tribes, respectively) and increased levels of speciation (Píchová *et al.* 2018), they represent ideal systems to test this hypothesis. It is postulated that sect. *Pusillae* was transferred to Africa (ca 50.3 Mya), while sect. *Claviceps* originated in North America (ca. 20.7 Mya), and it is likely that the common ancestor shared between these sections (Fig. 1) had strains that were transferred to Africa, likely due to insect vectors via transatlantic long-distance dispersal (Píchová *et al.* 2018). The strains that remained, in South America, likely persisted but appeared to not speciate for roughly 30 Mya (Píchová *et al.* 2018), despite having aspects of a more adaptable two-speed genome (Fig. 4, 5). Limited sampling records could be a factor contributing to this lack of speciation during this 30 My period, but it could also be suggested that the ancestral species of sects. *Claviceps* did not diverge due to a lack of diversification of host species (Píchová *et al.* 2018). It is well known that *Claviceps* species share a rather unique relationship with their hosts (strict ovarian parasites). The evolution of *Claviceps* appears to be primarily driven by the evolution and diversification of the host species (Píchová *et al.* 2018). This can be inferred from divergence time estimates which show that the crown node of sect. *Pusillae* aligns with the crown node of PACMAD grasses (ca. 45 Mya) (Bouchenak-Khelladi *et al.* 2010, Píchová *et al.* 2018), suggesting that these two organisms radiated in tandem after ancestral strains of sect. *Pusillae* were transferred to Africa. Similarly, the estimated crown node of sect. *Claviceps* corresponds with the origin of the core Pooideae (Poeae, Triticeae, Bromeae, and Littledaleae), which occurred in North America (ca. 33-26 Mya) (Bouchenak-Khelladi *et al.* 2010; Sandve & Fjellheim 2010).

Such a large difference between the estimate divergence age (~30 My) and long divergence branch (Fig. 1) between sect. *Clavcieps* and the other three sections (Píchová *et al.* 2018) suggests that a sudden event sparked the adaptive radiation within this section (Fig. 1). Under an assumption that ancestral strains of sect. *Claviceps* were infecting sedges (Cyperaceae), as is seen in the basal species *C. cyperi*, a host jump to BOP grasses could have ignited the rapid speciation of sect. *Claviceps*, similar to the suggested tandem radiation of sect. *Pusillae* with the PACMAD grasses in Africa. However, unknown factors might be responsible for the drastic genomic changes (i.e. putative loss of RIP) observed in sect. *Claviceps*, as no such changes were observed in sect. *Pusillae*. The radiation of the core Pooideae occurred after a global super-cooling period (ca. 33-26 Mya) in North America. During this period, Pooideae experienced a stress response gene family expansion which enabled adaptation and diversification to cooler, more open, habitats (Kellogg 2001; Sandve & Fjellheim 2010). As gene cluster expansion was observed in sect. *Claviceps* (the only section to infect BOP grasses) it suggests that the same environmental factors that caused the radiation of Pooideae could have similarly affected sect. *Claviceps* (Kondrashov 2012) and might have resulted in the host jump to Pooideae, and potentially other BOP tribes. Interestingly, one of the orthogroups significantly expanded in sect. *Claviceps* (OG0000016) contains proteins associated with a cold-adapted (Alias *et al.* 2014) serine peptidase S8 subtilase (MER0047718; S08.139) (Additional file 2: Table S8). Although the crown node of sect. *Claviceps* is estimated at approximately 5-10 My before the radiation of the core Pooideae, the 95% highest posterior density determined in Píchová *et al.* (2018) could indicate both radiation events occurred at similar times.

Further examination of *Claviceps* species in South and Central America needs to be conducted to better elucidate the evolution and dispersal of the genus (Píchová *et al.* 2018). Efforts should focus on the elusive *C. junci*, a pathogen of Juncaceae (rushes), which is thought to reside in sect. *Claviceps* based on morphological and geographic characteristics (Langdon 1952; Píchová *et al.* 2018). This species, and potentially others, will provide further insight into the early evolution of sect. *Claviceps* and could bridge the current gap between the environmental factors that sparked the radiation of the core Pooideae and sect. *Claviceps*.

## Acknowledgements

We thank Dr. Miroslav Kolařík for providing *Claviceps* isolates from the Culture Collection of Clavicipitaceae at Institute of Microbiology, Academy of Sciences of the Czech Republic (CCC samples); Parivash Shoukouhi, Dr. Jim Menzies, and Zlatko Popovic for collection, isolation, maintenance, and DNA extraction of LM samples; Dr. Chris Schardl and Dr. Neil Moore, University of Kentucky, for providing the 2013 GFF3 files for *C. paspali* and *C. fusiformis*; Dr. Joshua Weitz and the Franklin Graybill Statistical Laboratory at Colorado State University for their help in data analysis of genomic fluidity; Molecular Technologies Laboratory (MTL) at the Ottawa Research & Development Centre, Agriculture and Agri-Food Canada, especially Kasia Dadej for technical assistance. For genomes downloaded from JGI these sequence data were produced by the US Department of Energy Joint Genome Institute https://www.jgi.doe.gov/ in collaboration with the user community.

## Data availability

Datasets and scripts are available on Dryad: Wyka, Stephen *et al.* (2020), Genus-wide comparison reveals divergence and evolution of the four sections within the genus Claviceps are the result of varying mechanisms driving genome evolution and host range, Dryad, Dataset, https://doi.org/10.5061/dryad.18931zcsk. (Submitted upon publication)

Genomes and Illumina raw reads were deposited to NCBI under the BioProject PRJNA528707 (Additional file 1: Table S1). Scripts are maintained within the GitHub repository of the primary author’s, https://github.com/PlantDr430/CSU_scripts. TransposableELMT can be found at Zenodo doi: 105281/zenodo3469661. All phylogenetic trees were made available at TreeBase (ID:XXXX).

## Author Contributions

The project was conceived and designed by S.A.W., S.J.M., and K.B.; S.A.W. performed the research, annotations, bioinformatic workflows, and analyzed the data with technical troubleshooting from S.J.M.; M.L. and J.R.D initiated whole-genome sequencing of LM samples; M.L., V.N., and K.B. provided management, research advice, and editorial contributions; S.A.W. wrote the paper with contributions from all other authors.

