## Additional_File_1_Fig_S1-S14_Table_S1-S6 for "Whole genome comparisons of ergot fungi reveals the divergence and evolution of species within the genus *Claviceps* are the result of varying mechanisms driving genome evolution and host range expansion"

**Table of Contents:**


| | | |
|---|---|---|
| **Figure S1** | Page 2 | Mean functional proteins per sections |
| **Figure S2** | Page 3 | Neighbor-joining super-matrix phylogeny |
| **Figure S3** | Page 4 | Maximum parsimony super-matrix phylogeny |
| **Figure S4** | Page 5 | Density consensus of gene trees |
| **Figure S5** | Page 6 | *Claviceps* genus tree topology frequency |
| **Figure S6** | Page 7 | *Claviceps* section *Claviceps* tree topology frequency |
| **Figure S7** | Page 8 | *Claviceps* section *Pusillae* tree topology frequency |
| **Figure S8** | Page 9 | Transposable elements divergence landscapes |
| **Figure S9** | Page 10 | Distance of genes to closest transposable elements |
| **Figure S10** | Page 11 | Hexbin plots of intergenic flanking regions |
| **Figure S11** | Page 12 | Mean number of paralogs per orthogroups across sections |
| **Figure S12** | Page 13 | Heatmap of orthogroups with conserved domains |
| **Figure S13** | Page 14 | Heatmap of unclassified orthogroups |
| **Figure S14** | Page 15 | Number of orthogroups with expansion per section |
| **Table S1** | Page 16 | Collection and accession information of isolates |
| **Table S2** | Page 17 | Functional protein numbers per isolate |
| **Table S3** | Page 18 | Additional genomes used in OrthoFinder analysis |
| **Table S4** | Page 19 | *P*-values from genomic fluidity analysis |
| **Table S5** | Page 20 | Number of duplicated gene and percent in tandem |
| **Table S6** | Page 21 | RIP-index results from The RIPper |


**Additional files not located in this file.**

**Additional File 2:**

| | |
|---|---|
| **Table S7** | BLASTp results of 55 important *Claviceps* genes, including *rid-1* homolog |
| **Table S8** | Protein domain association of highly expanded orthogroups |
| **Table S9** | Order of orthogroups displayed in heatmaps |

**Additional File 3:**

| | |
|---|---|
| **Table S10-1** | All orthogroups with corresponding size, strains, gene IDs, and functional proteins |
| **Table S10-1** | classification, broken up into two files due to size. |

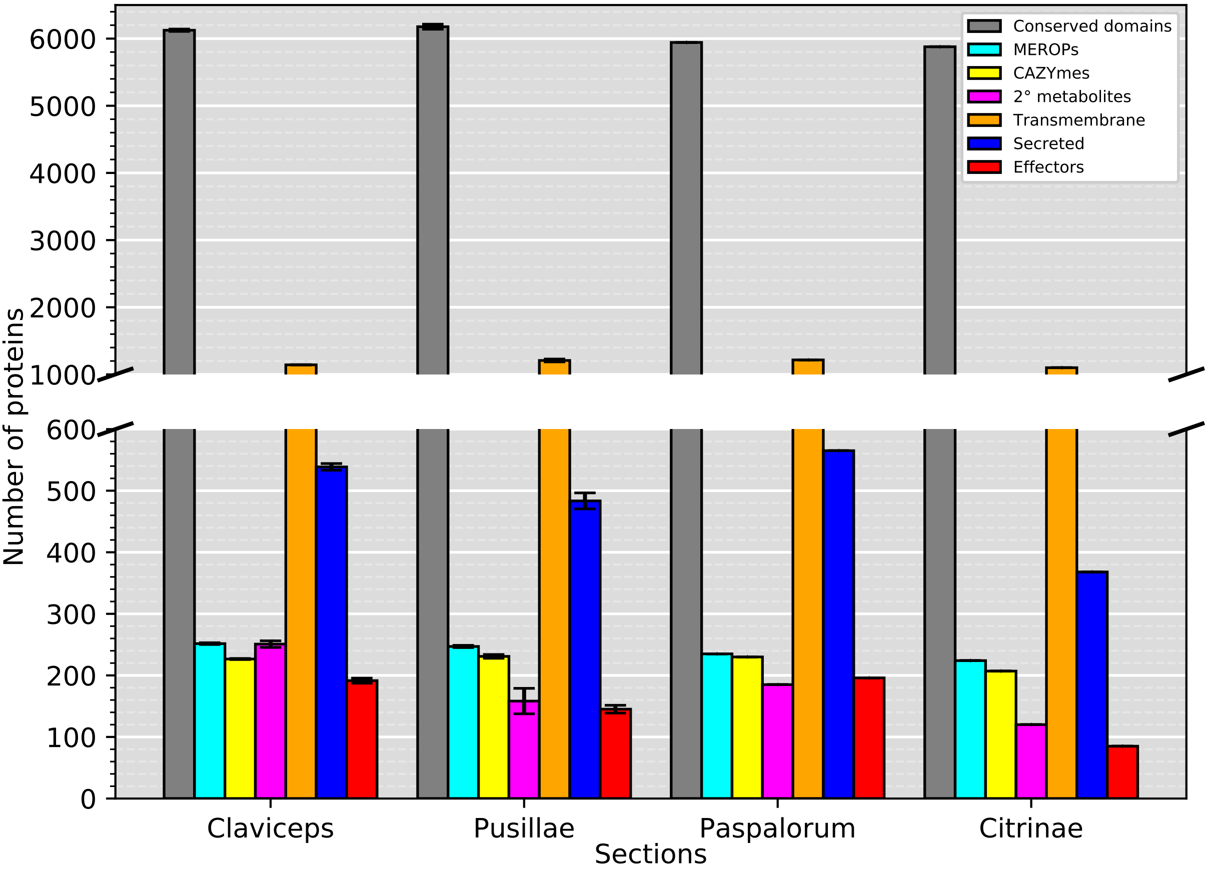


**Figure S1:** Mean number of proteins in each section of the genus *Claviceps*. Bars represent standard error.

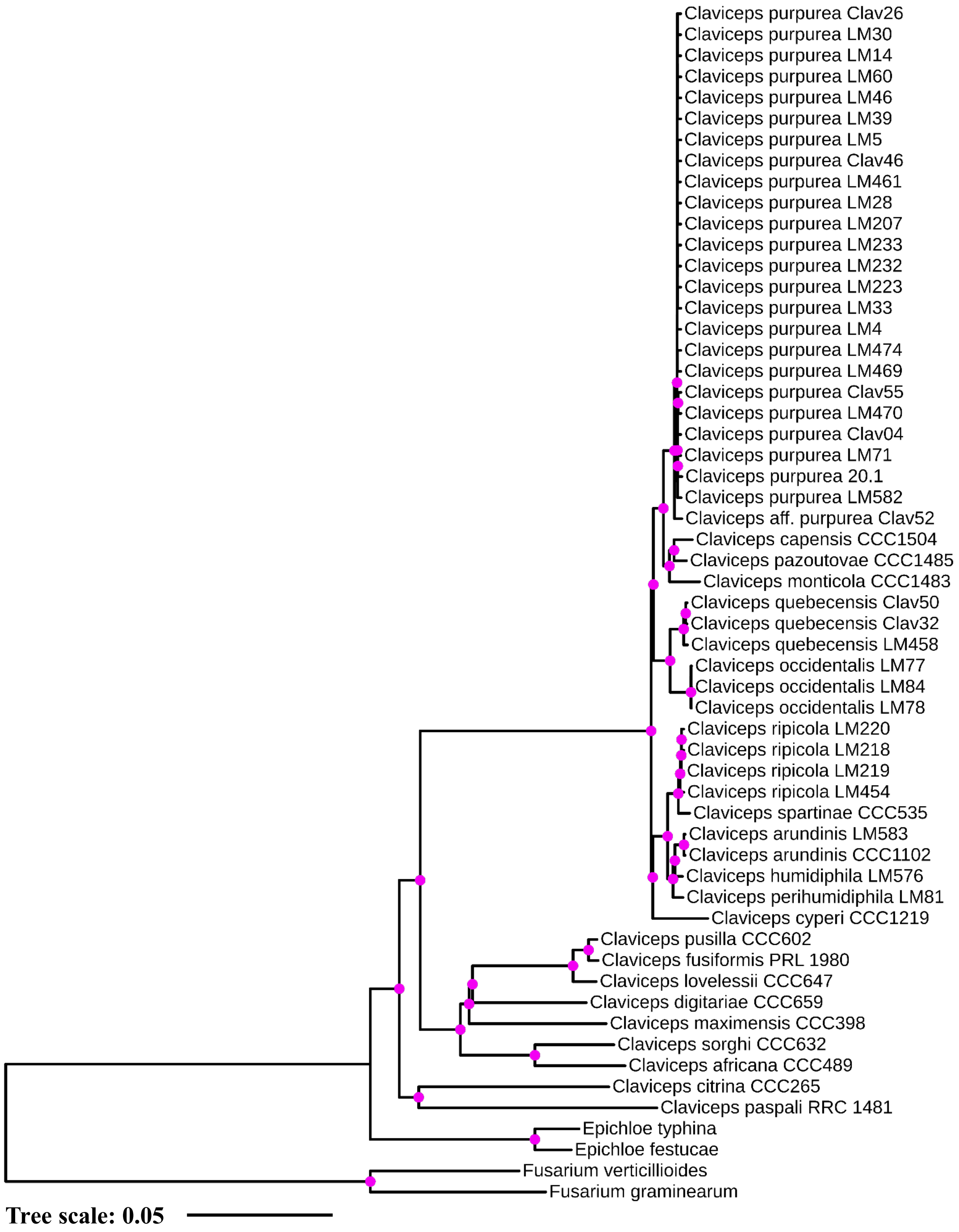


**Figure S2::** Neighbor- oining phylogenetic reconstruction of the *Claviceps* genus using amino acid sequences of 2,002 single copy orthologs with 1000 bootstrap replicates. Pin   dots at branches represent bootstrap values ≥95.

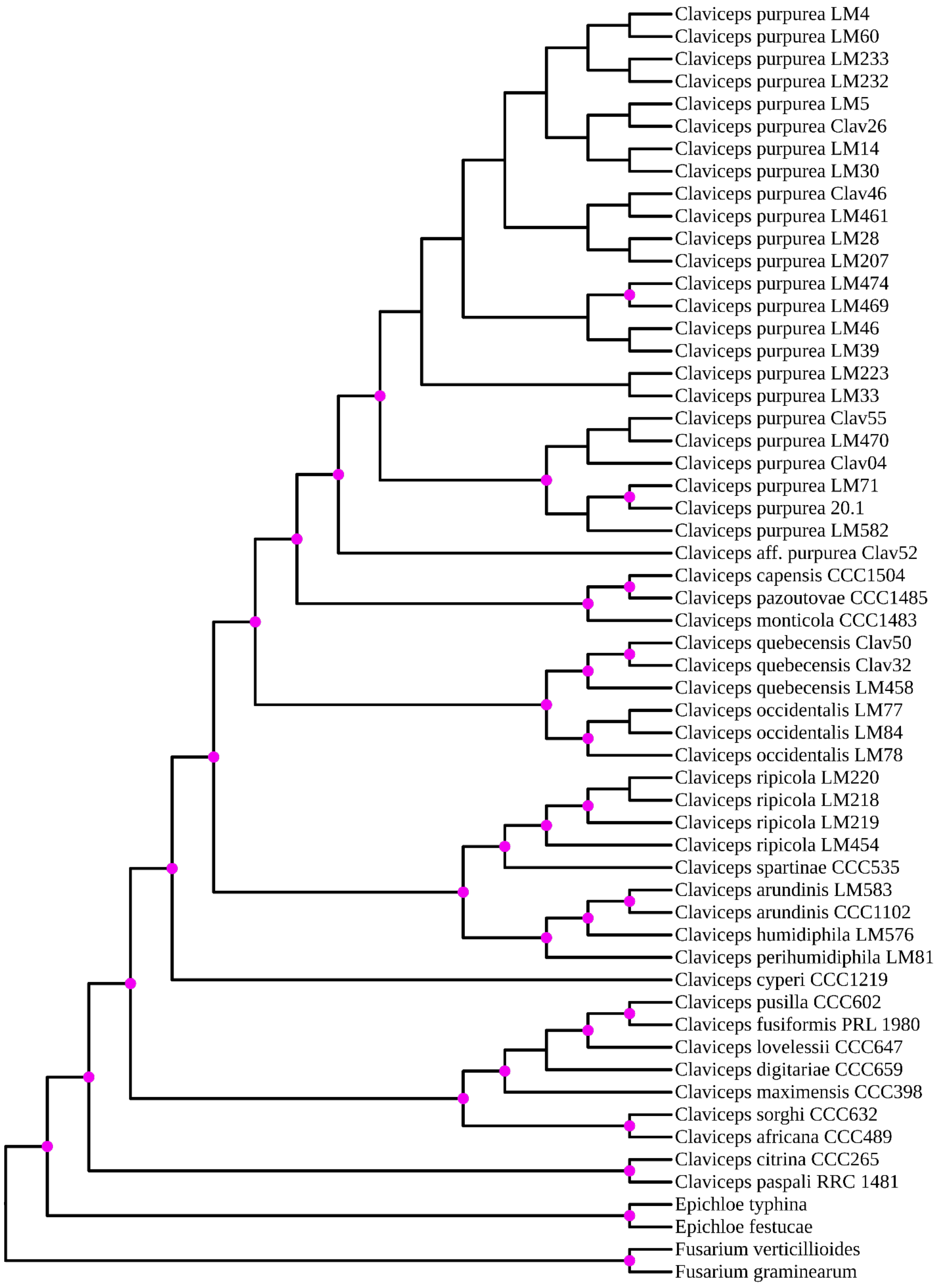


**Figure S3:** Maximum parsimony phylogenetic reconstruction of the *Claviceps* genus using amino acid sequences of 2,002 single copy orthologs with 1000 bootstrap replicates. Pin dots at branches represent bootstrap values ≥95.

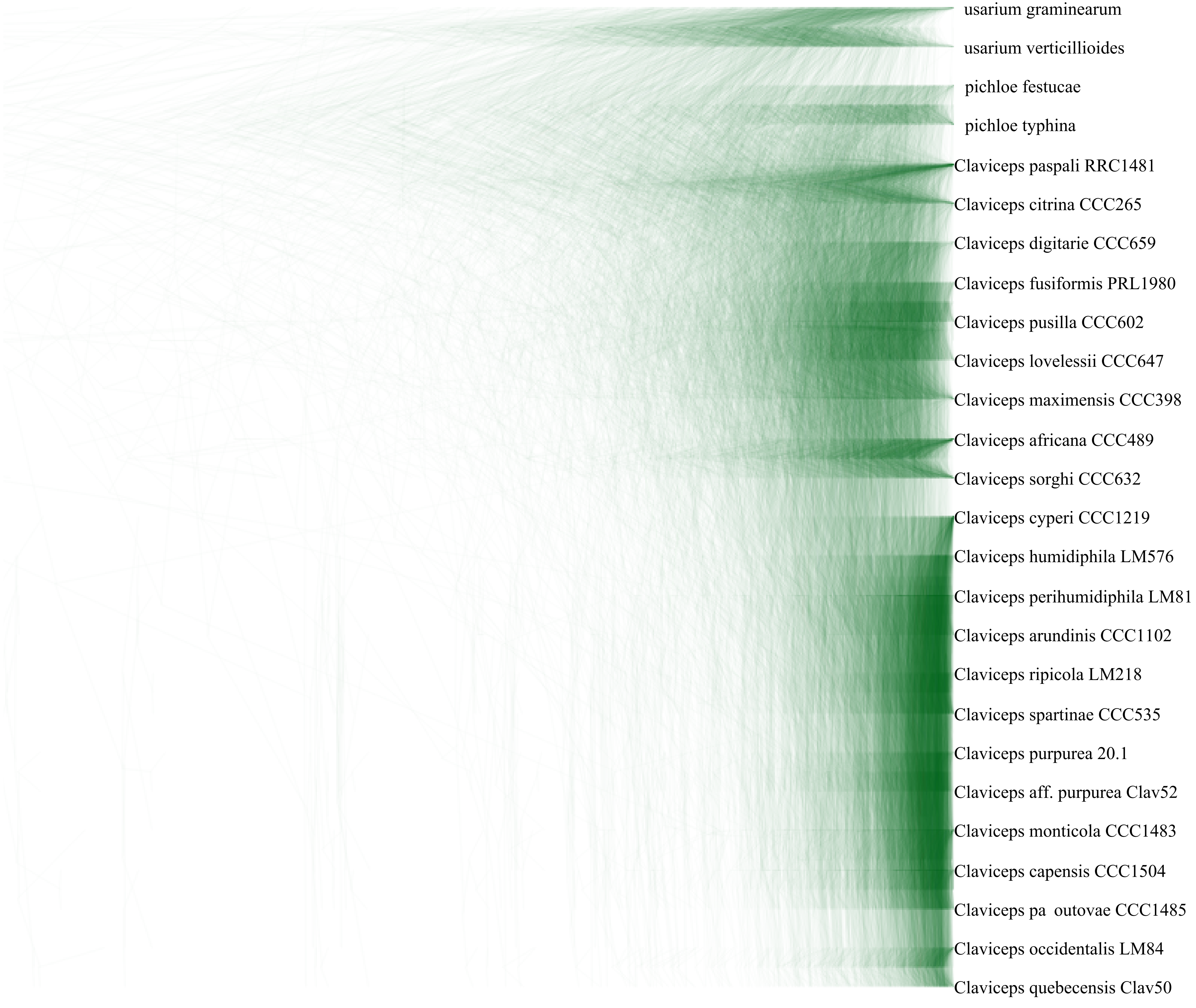


**Figure S4:** ensity consensus tree of 2,002 maximum li elihood phylogenetic reconstructions of the *Claviceps* genus using amino acid sequences of the single-copy orthologs with 1000 bootstrap replicates. Representative isolates from each species were used in this analysis for clarity. Thic er overlapping regions is an indicator of branch support. Tree order was determined by the most frequently occurring tree order.

**Figure S5:** Phylogenetic reconstructions and genealogy variation of gene trees for the *Claviceps* genus (excluding outgroups). (Line chart) Cumulative distribution of the number of genes per topology.    ori  ontal dotted lines indicate the half of the genes examined and total genes examined. (Trees)    our most frequent topologies with their corresponding frequencies.

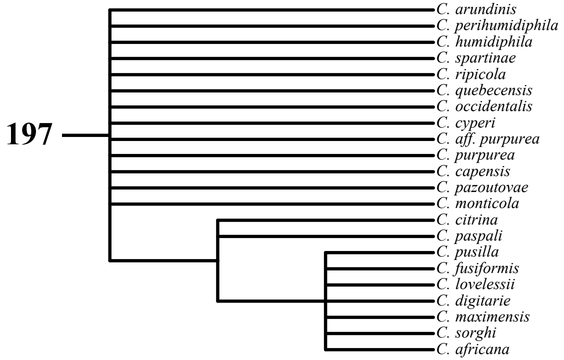


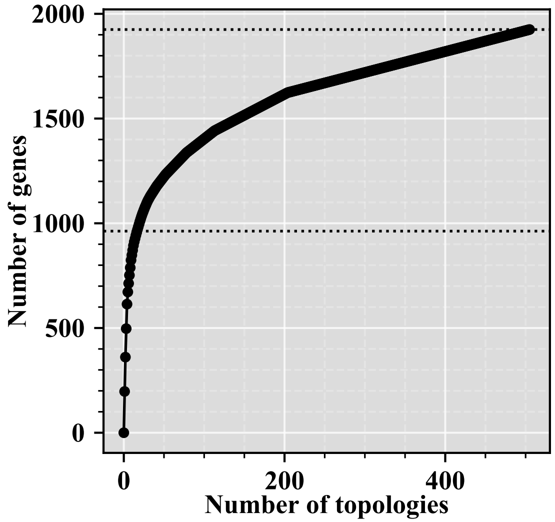


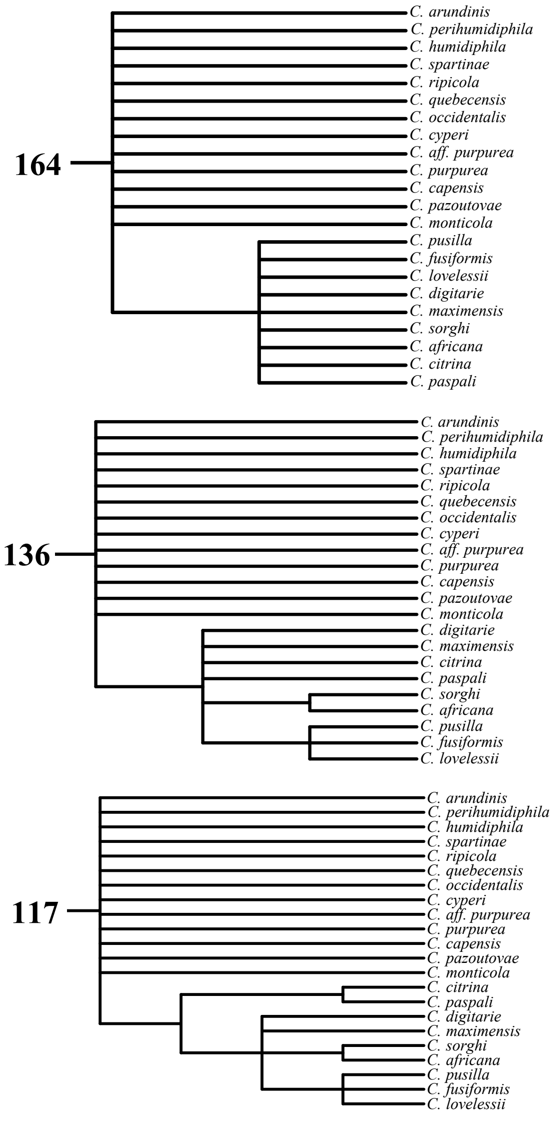

**Figure S6:** Phylogenetic reconstructions and genealogy variation of gene trees for *Claviceps* section *Claviceps*. (Line chart) Cumulative distribution of the number of genes per topology. ori ontal dotted lines indicate the half of the genes examined and total genes examined. (Trees) ix most frequent topologies with their corresponding frequencies.

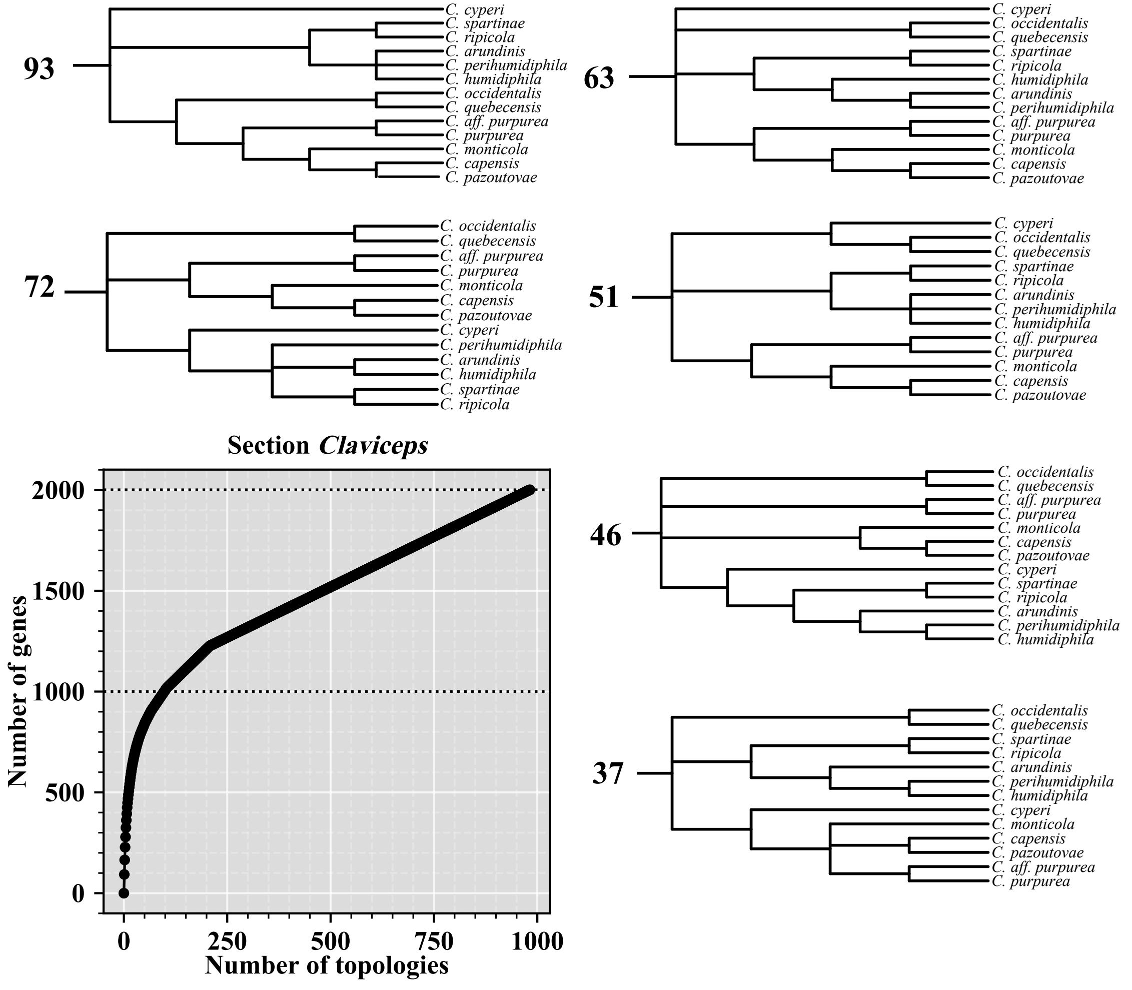

**Figure S7:** Phylogenetic reconstructions and genealogy variation of gene trees for *Claviceps* section *Pusillae*. (Line chart) Cumulative distribution of the number of genes per topology. ori ontal dotted lines indicate the half of the genes examined and total genes examined. (Trees) ix most frequent topologies with their corresponding frequencies.

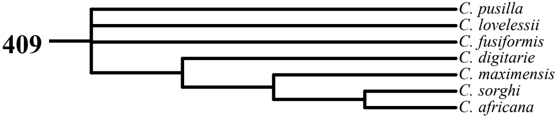


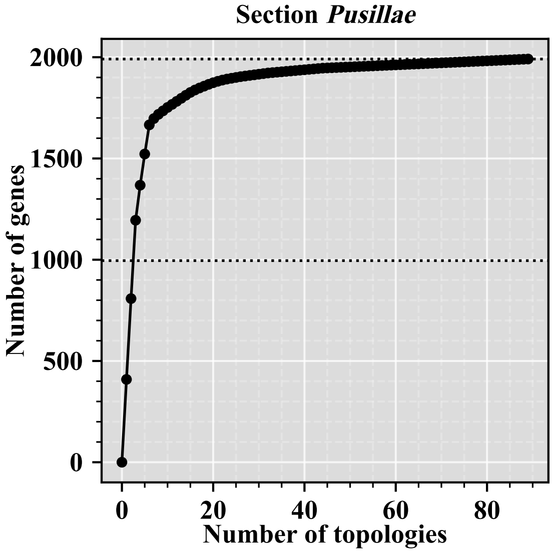


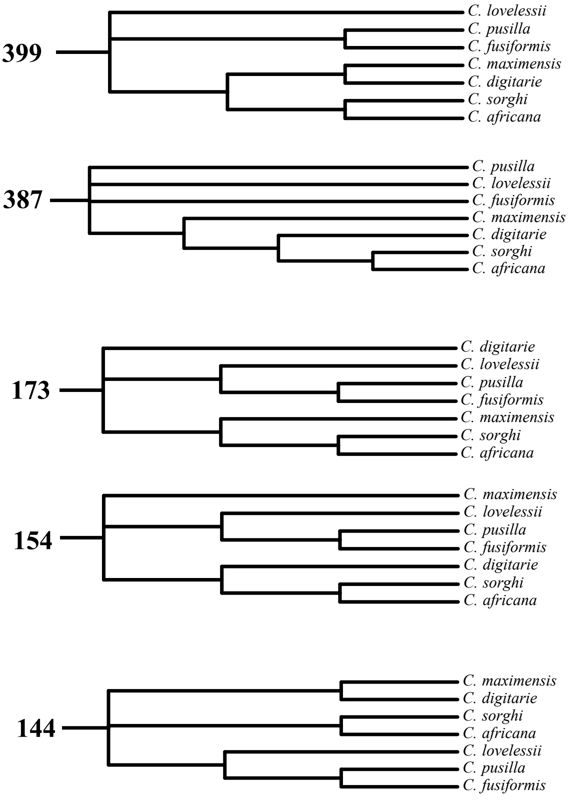

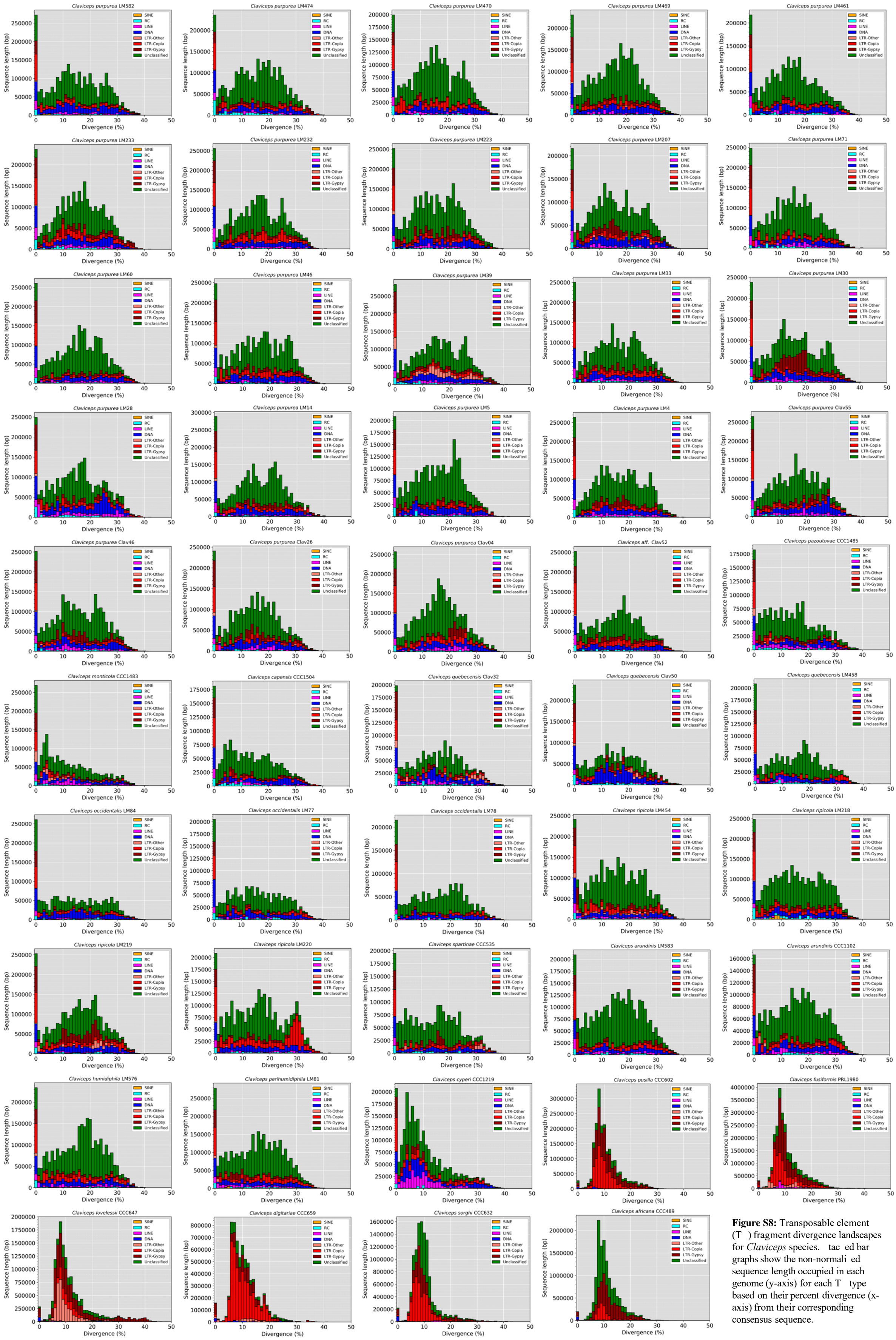


**Figure S8:** Transposable element (T ) fragment divergence landscapes for *Claviceps* species. tac ed bar graphs show the non-normali ed sequence length occupied in each genome (y-axis) for each T type based on their percent divergence (x-axis) from their corresponding consensus sequence.

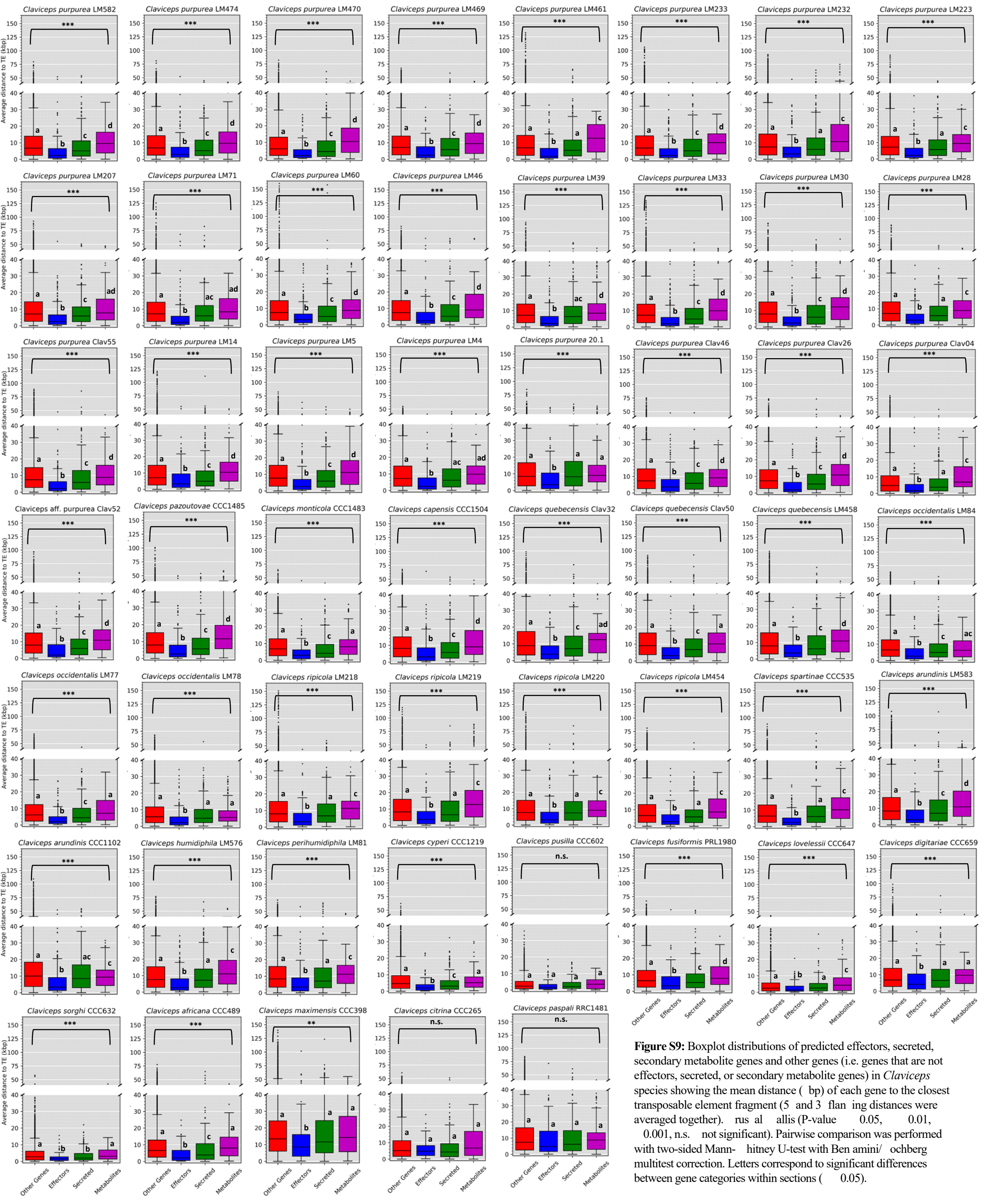


**Figure S9:** Boxplot distributions of predicted effectors, secreted, secondary metabolite genes and other genes (i.e. genes that are not effectors, secreted, or secondary metabolite genes) in *Claviceps* species showing the mean distance ( bp) of each gene to the closest transposable element fragment (5 and 3 flan ing distances were averaged together). rus al allis (P-value 0.05, 0.01, 0.001, n.s. not significant). Pairwise comparison was performed with two-sided Mann- hitney U-test with Ben amini/ ochberg multitest correction. Letters correspond to significant differences between gene categories within sections ( 0.05).

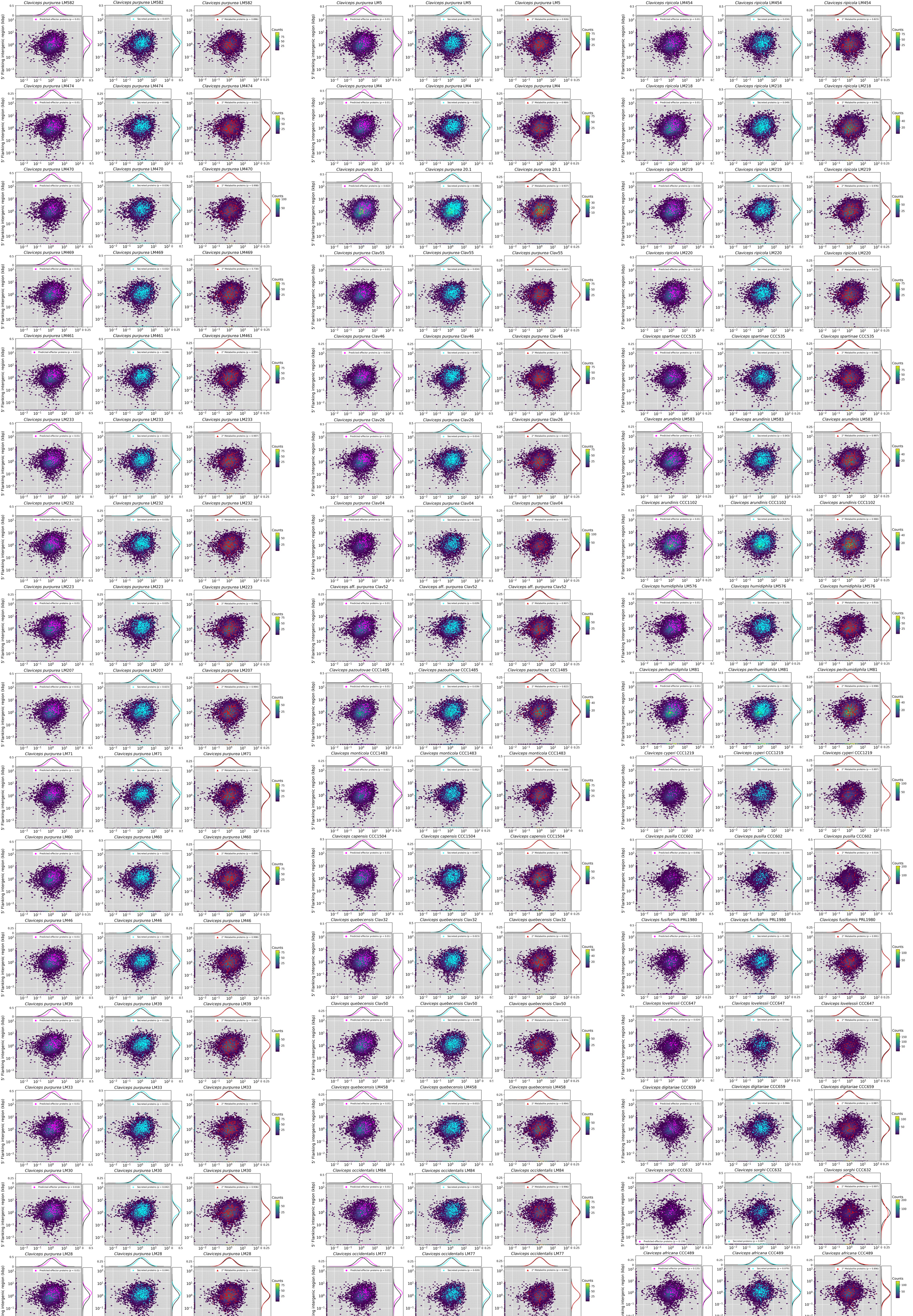


**Figure S10:** 5 and 3 intergenic region si e (y and x-axis) of *Claviceps* species. irst genes of contigs are plotted along the bottom of the x-axis and genes at the end of each contig are plotted along the y-axis. Colored hexbins indicate the intergenic lengths of all genes with color-code indicating the frequency distribution (gene counts) according to the legend on the right. verlaid mar ers indicate specific gene types corresponding to legends in the top right within each plot. requency distributions of specific gene types (corresponding legend color) and all other genes not of the specific type (blac ) are plotted along the x- and y-axis. or information on statistical test ee Methods.

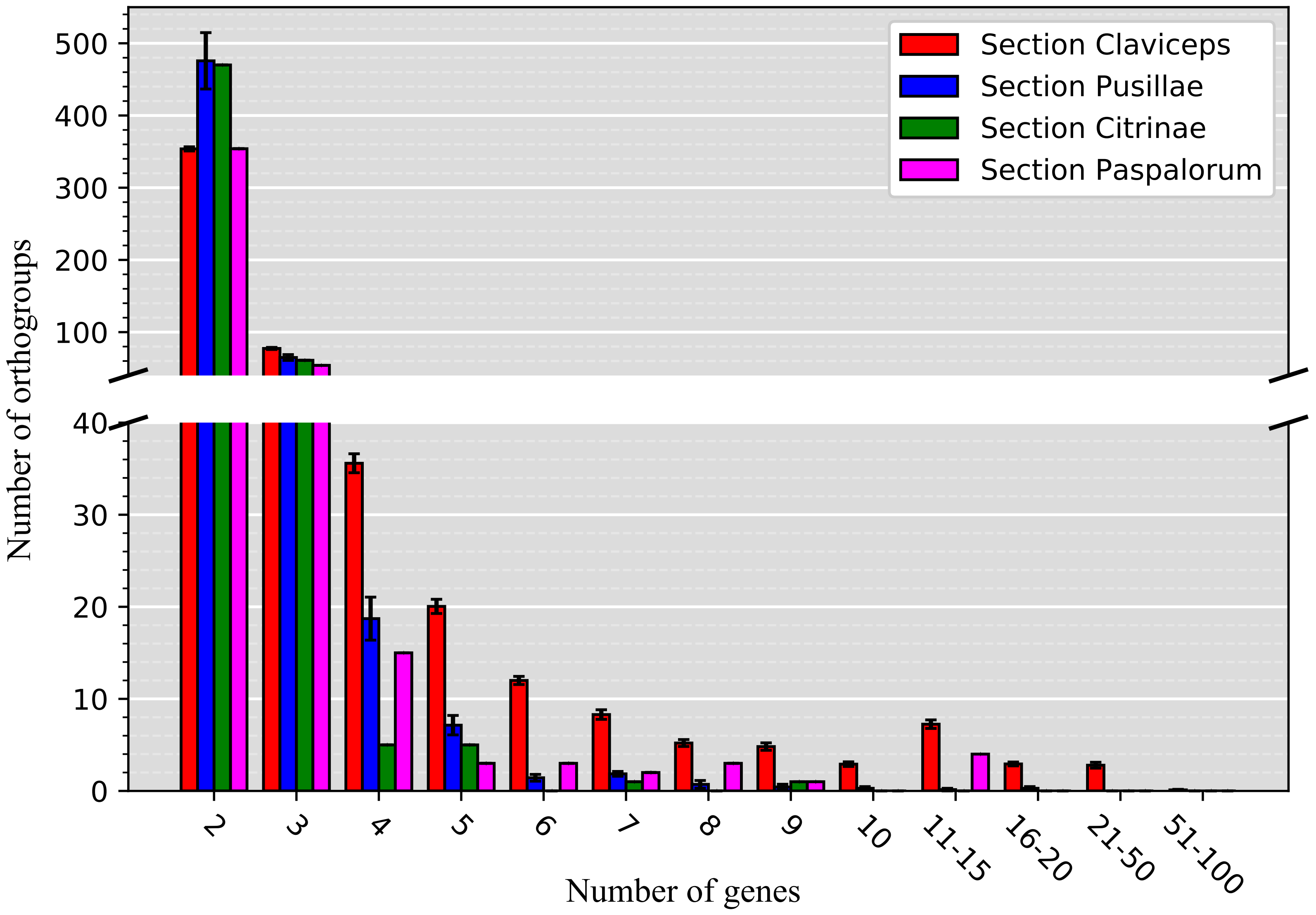


**Figure S11:** Mean number of orthogroups (y-axis) in each section of the genus Claviceps containing number of genes (x-axis), not including single gene orthogroups for better visuali ation of paralogs. Bars represent standard error.

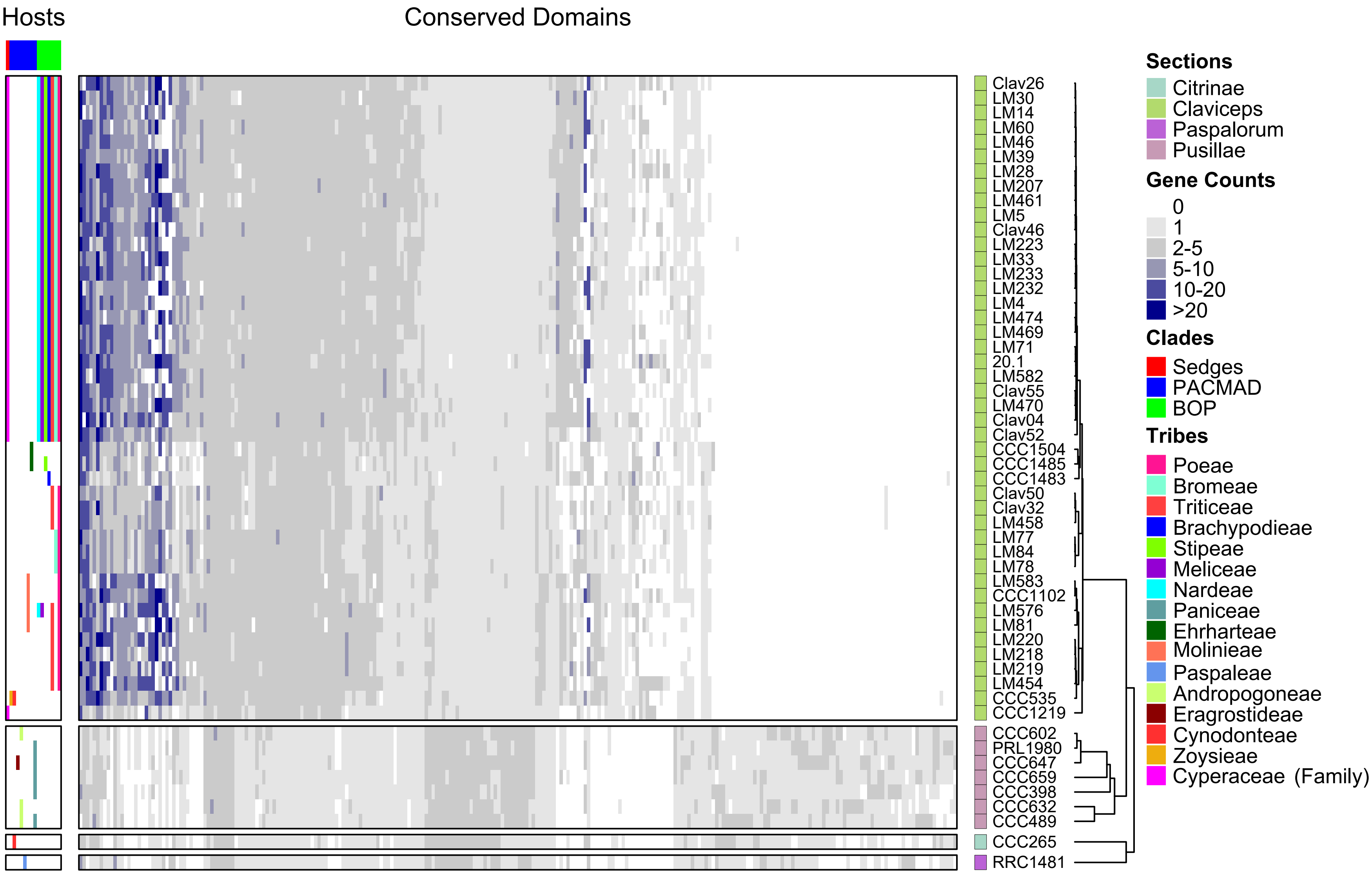


**Figure S12:** eatmap of gene counts in the remaining orthogroups containing genes encoding conserved protein domains for all 53 *Claviceps* strains ordered based on ML tree in ig. 1 and separated by sections. rthogroups are ordered based on hierarchical clustering. The host spectrum (left) is generali ed across species, as no literature has determined the existence of race specific isolates within species, is shown on the left side of the figure determined from literature review of field collected samples ( upplementary Material in P chov *et al.* 2018) and previous inoculation tests Campbell (1957) and Liu *et al.* (*Submitted*).

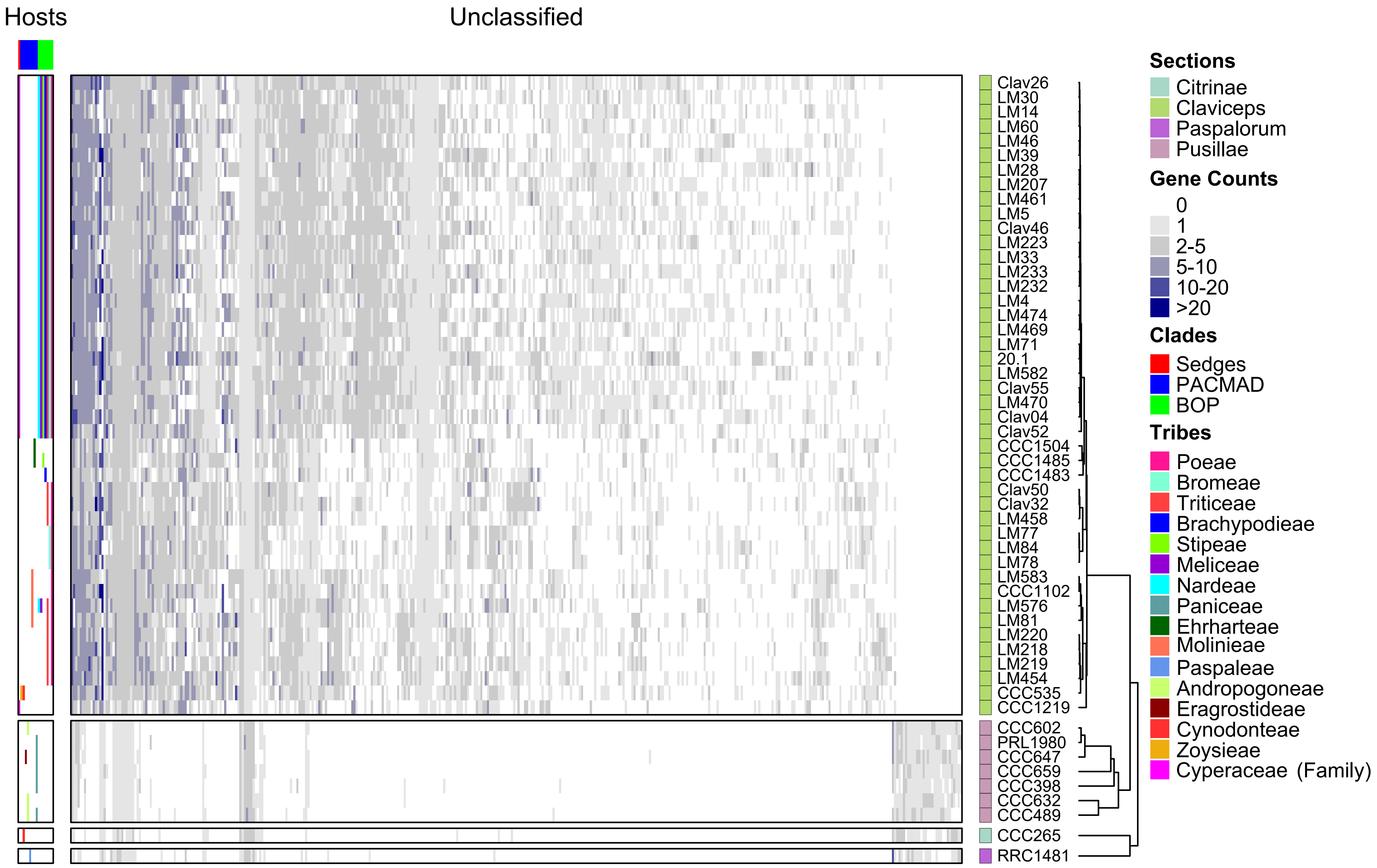


**Figure S13:** eatmap of gene counts in remaining orthogroups containing unclassified genes for all 53 *Claviceps* strains ordered based on ML tree in ig. 1 and separated by sections. rthogroups are ordered based on hierarchical clustering. The host spectrum (left) is generali ed across species, as no literature has determined the existence of race specific isolates within species, is shown on the left side of the figure determined from literature review of field collected samples ( upplementary Material in P chov *et al.* 2018) and previous inoculation tests Campbell (1957) and Liu *et al.* (*Submitted*).

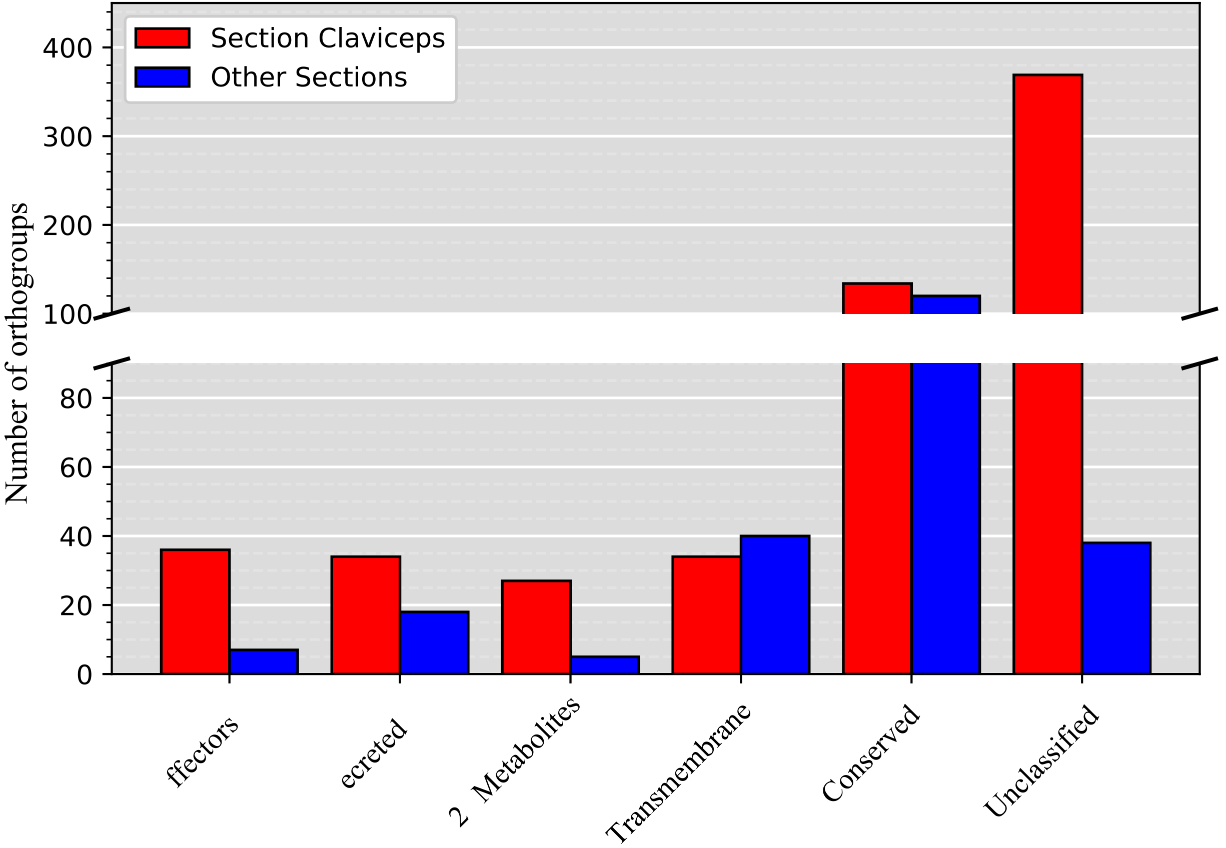


**Figure S14:** Number of orthogroups showing significantly ($P$ 0.01) greater expansion in respective *Claviceps* sections. ther sections include the combination of sects. *Pusillae*, *Citrinae*, and *Paspalorum*.

**Table S1:** Collection and accession information for strains used in this study.

| Organism | Strain | Strain alias | NCBI Accession | SRA Accession | Culture Collection | Location | Host | Collection Date |
|---|---|---|---|---|---|---|---|---|
| References: | | | | | | | | |
| *Claviceps purpruea* | 20.1 |  | AM A2272775 | -- | -- | Germany | *Secale cereale* | 1988 |
| *Claviceps fusiformis* | PRL 1980 | | AMN02981339 | -- | -- | Africa: Cote d Ivoire | *Pennisetum typhoideum* | 1958 |
| *Claviceps paspali* | RRC 1481 |  | AMN02981342 | -- | -- | U A: Georgia, Mansfield | *Paspalum* sp. | 2001 |
| This study: | | | | | | | | |
| *Claviceps purpruea* | Clav04 |  | AMN11159846 | RR8785178 |  | U A: Colorado, an Luis alley | *Bromus inermis* | 2016 |
| *Claviceps purpruea* | Clav26 |  | AMN11159847 | RR8785181 |  | U A: Colorado, an Luis alley | *Hordeum vulgare* | 2016 |
| *Claviceps purpruea* | Clav46 |  | AMN11159848 | RR8785180 |  | U A: yoming, orland | *Secale cereale* | 2016 |
| *Claviceps purpruea* | Clav55 |  | AMN11159850 | RR8785174 |  | New ealand | *Lolium perenne* | 2017 |
| *Claviceps purpruea* | LM4 |  | AMN11159851 | RR8785145 | A MC:250624 | Canada: Manitoba | *Tricosecale* | 1996 |
| *Claviceps purpruea* | LM5 |  | AMN11159852 | RR8785146 | A MC:250625 | Canada: Manitoba | *Hordeum vulgare* | 1996 |
| *Claviceps purpruea* | LM14 |  | AMN11159853 | RR8785147 | A MC:250634 | Canada: as atchewan | *Hordeum vulgare* | 1996 |
| *Claviceps purpruea* | LM28 | men ies 81 | AMN11159854 | RR6985966† | A MC:250647 | Canada: as atchewan | *Triticum aestivum* | 2000 |
| *Claviceps purpruea* | LM30 |  | AMN11159855 | RR8785151 | A MC:250649 | Canada: as atchewan | *Secale cereale* | 2000 |
| *Claviceps purpruea* | LM33 |  | AMN11159856 | RR8785141 | A MC:250652 | Canada: Manitoba | *Secale cereale* | 2015 |
| *Claviceps purpruea* | LM39 |  | AMN11159857 | RR8785142 | A MC:250658 | Canada: as atchewan | *Triticum turgidum subsp. durum* | 2000 |
| *Claviceps purpruea* | LM46 |  | AMN11159858 | RR8785143 | A MC:250663 | Canada: Alberta | *Triticum turgidum subsp. durum* | 2000 |
| *Claviceps purpruea* | LM60 |  | AMN11159859 | RR8785144 | A MC:250680 | Canada: Manitoba | *Avena sativa* | 2005 |
| *Claviceps purpruea* | LM71 |  | AMN11159860 | RR8785148 | A MC:250720 | United ingdom | *Alopercurus myosuroides* | 2004 |
| *Claviceps purpruea* | LM207 | | AMN11159861 | RR8785149 | | Canada: Manitoba | *Elymus repens* | 2014 |
| *Claviceps purpruea* | LM223 |  | AMN11159862 | RR8785164 | A MC:250814 | Canada: Manitoba | *Bromus riparius* | 2014 |
| *Claviceps purpruea* | LM232 |  | AMN11159863 | RR8785161 | A MC:250822 | Canada: Manitoba | *Phalaris canariensis* | 2014 |
| *Claviceps purpruea* | LM233 | | AMN11159864 | RR8785162 | | Canada: Manitoba | *Phalaris canariensis* | 2014 |
| *Claviceps purpruea* | LM461 |  | AMN11159865 | RR8785163 | A MC:251847 | Canada: uebec | *Elymus repens* | 2016 |
| *Claviceps purpruea* | LM469 |  | AMN11159866 | RR8785165 |  | Canada: ntario | *Triticum aestivum* | 2016 |
| *Claviceps purpruea* | LM470 |  | AMN11159867 | RR8785166 |  | Canada: ntario | *Elymus repens* | 2016 |
| *Claviceps purpruea* | LM474 |  | AMN11159868 | RR8785167 |  | Canada: ntario | *Hordeum vulgare* | 2016 |
| *Claviceps purpruea* | LM582 | CCC771 | AMN11159869 | RR6985962† | A MC:251723 | C ech Republic: Be dedice | *Secale cereale* | 2003 |
| *Claviceps aff. purpruea* | Clav52 |  | AMN11159849 | RR8785175 |  | U A: ashington | *Poa pratensis* | 2017 |
| *Claviceps quebecensis* | Clav32 |  | AMN11159882 | RR8785176 |  | U A: Montana, hephard | *Hordeum vulgare* | 2016 |
| *Claviceps quebecensis* | Clav50 |  | AMN11159881 | RR8785177 |  | U A: lahoma, oop house Ardmore | *Elymus sp.* | 2017 |
| *Claviceps quebecensis* | LM458 |  | AMN11159883 | RR6985957† | A MC:251898 | Canada: uebec, Cote Nord | *Ammophila* (plant) | 2015 |
| *Claviceps occidentalis* | LM77 |  | AMN11159879 | RR8785179 | A MC:250577 | Canada: Alberta | *Phleum pratense* | 2016 |
| *Claviceps occidentalis* | LM78 |  | AMN11159878 | RR6985960† | A MC:250578 | Canada: Alberta, North tar | *Bromus inermis* | 1956 |
| *Claviceps occidentalis* | LM84 |  | AMN11159876 | RR8785170 | A MC:250590 | Canada: British Columbia | *Bromus inermis* | 2016 |
| *Claviceps ripicola* | LM218 | M 7.2 | AMN11159875 | RR6985964† | A MC:251843 | Canada: Manitoba, Grants ield nowfla e | *Phalaris arundinacea* | 2014 |
| *Claviceps ripicola* | LM219 |  | AMN11159874 | RR8785169 | A MC:250811 | Canada: Manitoba | *Phalaris arundinacea* | 2014 |
| *Claviceps ripicola* | LM220 |  | AMN11159873 | RR8785168 | A MC:250812 | Canada: Manitoba | *Phalaris arundinacea* | 2014 |
| *Claviceps ripicola* | LM454 | 139 | AMN11159872 | RR6985963† | A MC:251845 | Canada: uebec, MRC Maria-Chapdelaine | *Ammophila breviligulata* | 2014 |
| *Claviceps spartinae* | CCC535 |  | AMN11159888 | RR8785160 | CCC:535 | United ingdom: Marchwood | *Sporobolus anglicus* | 1999 |
| *Claviceps arundinis* | LM583 | CCC933 | AMN11159894 | RR6985961† | A MC:251724/CCC:933 | C ech Republic: a lovy vory, tary rbens y pond | *Phragmites australis* | 2008 |
| *Claviceps arundinis* | CCC1102 |  | AMN11159893 | RR8785153 | CCC:1102 | rance: 973 Rte de Beaune | *Phragmites australis* | 2009 |
| *Claviceps humidiphila* | LM576 | CCC434 | AMN11159871 | RR6985959† | A MC:251717/CCC:434 | Germany: Bavaria | *Dactylis sp.* | 1998 |
| *Claviceps perihumidiphila* | LM81 |  | AMN11159877 | RR6985958† | A MC:250581 | Canada: Alberta, Metis ow | *Elymus albicans* | 1956 |
| *Claviceps cyperi* | CCC1219 |  | AMN11159895 | RR8785154 | CCC:1219 | outh Africa: empton Par | *Cyperus esculentus* | 2012 |
| *Claviceps capensis* | CCC1504 |  | AMN11159898 | RR8785171 | CCC:1504T | outh Africa: Cape Town, estern Cape | *Ehrharta villosa* | 2014 |
| *Claviceps pazoutovae* | CCC1485 |  | AMN11159897 | RR8785152 | CCC:1485T | outh Africa: ogsbac , astern Cape | *Stipa dregeana* | 2014 |
| *Claviceps monticola* | CCC1483 |  | AMN11159896 | RR8785150 | CCC:1483T | outh Africa: ogsbac , astern Cape | *Brachypodium sp.* | 2014 |
| *Claviceps pusilla* | CCC602 | | AMN11159889 | RR8785157 | CCC:602 | imbabwe: Matopos | *Bothriochloa insculpta* | 2000 |
| *Claviceps lovelessii* | CCC647 |  | AMN11159891 | RR8785155 | CCC:647T | imbabwe: Matopos, Matopos Research tation | *Eragrostis sp.* | 2001 |
| *Claviceps digitariae* | CCC659 | | AMN11159892 | RR8785156 | CCC:659 | Africa: Botswana | *Digitaria eriantha* | -- |
| *Claviceps maximensis* | CCC398 | | AMN11159886 | RR8785172 | CCC:398 | Paraguay: Chaco | *Megathyrsus maximus* | 1997 |
| *Claviceps sorghi* | CCC632 |  | AMN11159890 | RR8785158 | CCC:632 | India: arnata a, ewargi, Gulbarga | *Sorghum bicolor* | 2000 |
| *Claviceps africana* | CCC489 | | AMN11159887 | RR8785159 | CCC:489 | Mexico: Celaya, Guana uato | *Sorghum bicolor* | 1998 |
| *Claviceps citrina* | CCC265 |  | AMN11159885 | RR8785173 | CCC:265 | Mexico: Texcoco (semillero) 6.5 m to Mexico City | *Distichlis spicata* | 1996 |

† RA data first published in Nguyen *et al.* 2018
Cultures available at the lab of  r.  amsi Nalam, Colorado  tate University,  ort Collins, C  or  r. Miao Liu  ttawa, Research and  evelopment Centre, Agriculture and Agri- ood Canada,  ttawa, Canada

**Table S2:** Number of genes with functional protein classifications for all 53 *Claviceps* genomes in this study.

| Organism | Strain | Section | Protein function | | | | | | |
|---|---|---|---|---|---|---|---|---|---|
| | | | Conserved domains | MEROP domains | CAZY -mes | 2° meta-bolites | Trans-membrane | Secreted signals | Predicted effectors |
| **References:** | | | | | | | | | |
| *C. purpruea* | 20.1 | Claviceps | 6560 | 255 | 243 | 321 | 1114 | 547 | 199 |
| *C. fusiformis* | PRL1980 | Pusillae | 6178 | 249 | 244 | 227 | 1315 | 521 | 152 |
| *C. paspali* | RRC1481 | Paspalorum | 5943 | 235 | 230 | 185 | 1216 | 565 | 196 |
| **This study:** | | | | | | | | | |
| *C. purpruea* | Clav04 | Claviceps | 6233 | 246 | 231 | 199 | 1145 | 557 | 221 |
| *C. purpruea* | Clav26 | Claviceps | 6221 | 263 | 229 | 255 | 1158 | 592 | 226 |
| *C. purpruea* | Clav46 | Claviceps | 6181 | 266 | 229 | 281 | 1151 | 560 | 198 |
| *C. purpruea* | Clav55 | Claviceps | 6113 | 260 | 231 | 253 | 1166 | 553 | 195 |
| *C. purpruea* | LM4 | Claviceps | 6132 | 249 | 233 | 230 | 1149 | 584 | 231 |
| *C. purpruea* | LM5 | Claviceps | 6179 | 257 | 231 | 287 | 1154 | 547 | 193 |
| *C. purpruea* | LM14 | Claviceps | 6132 | 259 | 227 | 273 | 1164 | 529 | 172 |
| *C. purpruea* | LM28 | Claviceps | 6260 | 254 | 233 | 279 | 1156 | 540 | 188 |
| *C. purpruea* | LM30 | Claviceps | 6167 | 262 | 229 | 268 | 1156 | 588 | 220 |
| *C. purpruea* | LM33 | Claviceps | 6193 | 263 | 226 | 264 | 1143 | 597 | 237 |
| *C. purpruea* | LM39 | Claviceps | 6156 | 265 | 232 | 256 | 1150 | 584 | 228 |
| *C. purpruea* | LM46 | Claviceps | 6125 | 261 | 225 | 250 | 1158 | 554 | 201 |
| *C. purpruea* | LM60 | Claviceps | 6126 | 256 | 231 | 290 | 1147 | 553 | 195 |
| *C. purpruea* | LM71 | Claviceps | 6095 | 240 | 221 | 248 | 1152 | 550 | 219 |
| *C. purpruea* | LM207 | Claviceps | 6120 | 259 | 220 | 251 | 1140 | 571 | 224 |
| *C. purpruea* | LM223 | Claviceps | 6076 | 234 | 230 | 250 | 1146 | 564 | 214 |
| *C. purpruea* | LM232 | Claviceps | 6169 | 260 | 232 | 284 | 1156 | 555 | 199 |
| *C. purpruea* | LM233 | Claviceps | 6294 | 264 | 230 | 266 | 1162 | 570 | 207 |
| *C. purpruea* | LM461 | Claviceps | 6223 | 237 | 231 | 246 | 1154 | 550 | 210 |
| *C. purpruea* | LM469 | Claviceps | 6106 | 255 | 229 | 273 | 1148 | 542 | 185 |
| *C. purpruea* | LM470 | Claviceps | 6195 | 256 | 223 | 210 | 1144 | 533 | 184 |
| *C. purpruea* | LM474 | Claviceps | 6118 | 242 | 222 | 290 | 1162 | 522 | 191 |
| *C. purpruea* | LM582 | Claviceps | 6132 | 257 | 226 | 229 | 1141 | 545 | 198 |
| *C. aff. purpruea* | Clav52 | Claviceps | 6078 | 252 | 225 | 250 | 1153 | 523 | 177 |
| *C. quebecensis* | Clav32 | Claviceps | 6057 | 244 | 226 | 266 | 1157 | 522 | 174 |
| *C. quebecensis* | Clav50 | Claviceps | 5986 | 248 | 228 | 260 | 1139 | 524 | 174 |
| *C. quebecensis* | LM458 | Claviceps | 6007 | 243 | 226 | 235 | 1135 | 508 | 154 |
| *C. occidentalis* | LM77 | Claviceps | 6020 | 243 | 222 | 186 | 1132 | 517 | 182 |
| *C. occidentalis* | LM78 | Claviceps | 6052 | 246 | 223 | 163 | 1133 | 513 | 174 |
| *C. occidentalis* | LM84 | Claviceps | 6088 | 244 | 223 | 189 | 1129 | 517 | 180 |
| *C. ripicola* | LM218 | Claviceps | 6090 | 249 | 228 | 255 | 1136 | 545 | 203 |
| *C. ripicola* | LM219 | Claviceps | 6133 | 255 | 226 | 270 | 1130 | 538 | 188 |
| *C. ripicola* | LM220 | Claviceps | 6168 | 249 | 225 | 255 | 1122 | 564 | 211 |
| *C. ripicola* | LM454 | Claviceps | 6163 | 261 | 224 | 261 | 1134 | 554 | 210 |
| *C. spartinae* | CCC535 | Claviceps | 6146 | 260 | 227 | 249 | 1153 | 504 | 163 |
| *C. arundinis* | LM583 | Claviceps | 6102 | 253 | 225 | 287 | 1140 | 526 | 186 |
| *C. arundinis* | CCC1102 | Claviceps | 6213 | 259 | 226 | 288 | 1130 | 548 | 195 |
| *C. humidiphila* | LM576 | Claviceps | 6147 | 253 | 227 | 282 | 1152 | 564 | 211 |
| *C. perihumidiphila* | LM81 | Claviceps | 6106 | 238 | 225 | 270 | 1124 | 522 | 176 |
| *C. cyperi* | CCC1219 | Claviceps | 5774 | 228 | 202 | 135 | 1072 | 392 | 97 |
| *C. capensis* | CCC1504 | Claviceps | 5989 | 245 | 225 | 250 | 1145 | 475 | 137 |
| *C. pazoutovae* | CCC1485 | Claviceps | 5954 | 232 | 223 | 225 | 1137 | 478 | 150 |
| *C. monticola* | CCC1483 | Claviceps | 5908 | 248 | 220 | 207 | 1118 | 481 | 148 |
| *C. pusilla* | CCC602 | Pusillae | 6276 | 252 | 227 | 76 | 1235 | 461 | 138 |
| *C. lovelessii* | CCC647 | Pusillae | 6351 | 248 | 228 | 110 | 1230 | 525 | 174 |
| *C. digitariae* | CCC659 | Pusillae | 6195 | 254 | 239 | 187 | 1213 | 513 | 158 |
| *C. maximensis* | CCC398 | Pusillae | 6100 | 244 | 235 | 226 | 1172 | 468 | 126 |
| *C. sorghi* | CCC632 | Pusillae | 6085 | 241 | 222 | 117 | 1139 | 425 | 123 |
| *C. africana* | CCC489 | Pusillae | 6057 | 241 | 221 | 165 | 1151 | 471 | 145 |
| *C. citrina* | CCC265 | Citrinae | 5879 | 224 | 207 | 120 | 1100 | 368 | 85 |

**Table S3:** Additional annotated genomes used in rtho inder analysis for finding orthologous gene clusters (orthogroups).

| **Organism** | **Strain** | **Accession** |
|---|---|---|
| *Acremonium chrysogenum* | ATCC 11550 | AMN02799700 |
| *Atkinsonella hypoxylon* | B4728 | http://www.endophyte.u y.edu/ |
| *Atkinsonella texensis* | B6155 | http://www.endophyte.u y.edu/ |
| *Balansia obtecta* | B249 | http://www.endophyte.u y.edu/ |
| *Clonostachys rosea* | CB 125111 | GI:1032557 |
| *Epichloe amarillians* | ATCC 200744 | http://www.endophyte.u y.edu/ |
| *Epichloe aotearoea* | ATCC M A-1229 | http://www.endophyte.u y.edu/ |
| *Epichloe baconii* | ATCC 200745 | http://www.endophyte.u y.edu/ |
| *Epichloe brachyelytri* | 4804 | http://www.endophyte.u y.edu/ |
| *Epichloe bromicola* | AL0426/2 | http://www.endophyte.u y.edu/ |
| *Epichloe coenophiala* | e4163 | http://www.endophyte.u y.edu/ |
| *Epichloe elymi* | ATCC 201551 | http://www.endophyte.u y.edu/ |
| *Epichloe festucae* | l1 | http://www.endophyte.u y.edu/ |
| *Epichloe gansuensis* | C M-2007b | http://www.endophyte.u y.edu/ |
| *Epichloe glyceriae* | ATCC 200747 | http://www.endophyte.u y.edu/ |
| *Epichloe inebrians* | ATCC M A-1228 | http://www.endophyte.u y.edu/ |
| *Epichloe mollis* | AL9924 | http://www.endophyte.u y.edu/ |
| *Epichloe sylvatica* | GR 10156 | http://www.endophyte.u y.edu/ |
| *Epichloe typhina* | ATCC 200736 | http://www.endophyte.u y.edu/ |
| *Epichloe uncinata* | CB 102646 | http://www.endophyte.u y.edu/ |
| *Fusarium ambrosium* | NRRL 20438 | AMN07200640 |
| *Fusarium avenaceum* | ave L 27 | AMN02850900 |
| *Fusarium fujikuroi* | | AM A4440726 |
| *Fusarium fujikuroi* | B14 | AM A4436914 |
| *Fusarium fujikuroi* | C1995 | AM A4440729 |
| *Fusarium fujikuroi* | 282 | AM A4440730 |
| *Fusarium fujikuroi* | G C8932 | AMN03075939 |
| *Fusarium fujikuroi* | U48 | AM A4440731 |
| *Fusarium fujikuroi* | IMI58289 | AM A3724789 |
| *Fusarium fujikuroi* | U3368 | AMN03075941 |
| *Fusarium fujikuroi* | U 10626 | AMN03075940 |
| *Fusarium fujikuroi* | m657 | AM A4440732 |
| *Fusarium fujikuroi* | MRC2276 | AM A4440733 |
| *Fusarium fujikuroi* | NCIM1100 | AM A4440734 |
| *Fusarium graminearum* | P -1/NRRL 31084 | AMN02953593 |
| *Fusarium kuroshium* | A -12 | AMN07200645 |
| *Fusarium langsethiae* | l201059 | AMN03274931 |
| *Fusarium longipes* | NRRL 20695 | AMN08631279 |
| *Fusarium mangiferae* | MRC7560 | AM A3862491 |
| *Fusarium oxysporum f. sp. cepae* | oC us2 | AMN05529097 |
| *Fusarium oxysporum f. sp. conglutinans* | 54008 | AMN02981380 |
| *Fusarium oxysporum f. sp. cubense* | 54006 | AMN02981379 |
| *Fusarium oxysporum f. sp. lycopersici* | 4287 | AMN02953675 |
| *Fusarium oxysporum f. sp. melonis* | 26406 | AMN02981378 |
| *Fusarium oxysporum f. sp. narcissi* | N139 | AMN05526391 |
| *Fusarium oxysporum f. sp. pisi* | 247 | AMN02981366 |
| *Fusarium oxysporum f. sp. radicis-cucumerinum* | orc016 | AMN04348764 |
| *Fusarium oxysporum f. sp. raphani* | 54005 | AMN02981381 |
| *Fusarium oxysporum f. sp. vasinfectum* | 25433 | AMN02981377 |
| *Fusarium poae* | 2516 | AMN05178635 |
| *Fusarium proliferatum* | T1 | AM A3862493 |
| *Fusarium pseudograminearum* | C 3096 | AMN02981337 |
| *Fusarium solani (Nectria haematococca)* | 77-13-4 | AMN02746079 |
| *Fusarium sporotrichioides* | NRRL 3299 | AMN08631227 |
| *Fusarium venenatum* | A3/5 | AM A2827224 |
| *Fusarium verticillioides* | 7600 | AMN02953630 |
| *Periglandula ipomoeae* | Iasa 13 | http://www.endophyte.u y.edu/ |
| *Purpureocillium lilacinum* | PL -1 | AMN04404347 |
| *Saccharomyces cerevisiae* | 288C | PR NA43747 |
| *Stachybotrys chlorohalonata* | IBT 40285 | AMN01819006 |
| *Trichoderma arundinaceum* | IBT 40837 | AMN06320351 |
| *Trichoderma asperellum* | CB 433.97 | AMN00769595 |
| *Trichoderma atroviride* | IMI 206040 | AMN02744066 |
| *Trichoderma citrinoviride* | TUCIM 6016 | AMN05369575 |
| *Trichoderma gamsii* | T6085 | AMN02849381 |
| *Trichoderma guizhouense* | N AU 4742 | AMN04535176 |
| *Trichoderma harzianum* | CB 22695 | AMN00761861 |
| *Trichoderma harzianum* | T6776 | AMN02851310 |
| *Trichoderma harzianum* | Tr1 | AMN06219536 |
| *Trichoderma harzianum* | TR274 | AMN07456232 |
| *Trichoderma harzianum* | M10 v1 | GI:1185309 |
| *Trichoderma harzianum* | T22 v1 | GI:1185313 |
| *Trichoderma longibrachiatum* | ATCC 18648 | AMN00767620 |
| *Trichoderma parareesei* | CB 125925 | AMN03784587 |
| *Trichoderma reesei* | M6a | AMN02746107 |
| *Trichoderma virens* | Gv29-8 | AMN02744059 |
| *Ustilaginoidea virens* | U -8b | AMN02693461 |
| *Ustilago maydis* | 521 | AMN02900459 |

**Table S4:** *P*-values for genomic fluidity differences from two-sample two-sided -test, bold numbers indicate significance.

| | usarium-Genus | pichloe-Genus | Trichoderma-Genus | T.hara ium-pecies | Pusillae-ection | Claviceps-Genus | .oxysporum-pecies | Claviceps-ection | C.purpurea-pecies | .fu i uroi-pecies |
|---|---|---|---|---|---|---|---|---|---|---|
| usarium-Genus | - | 8.74 -01 | 3.78 -01 | **3.30E-04** | **1.54E-03** | **7.98E-05** | **4.01E-07** | **4.35E-21** | **6.44E-27** | **3.65E-25** |
| pichloe-Genus | - | - | 4.94 -01 | **2.68E-03** | **6.47E-03** | **7.59E-04** | **4.39E-05** | **1.81E-14** | **2.04E-18** | **9.37E-18** |
| Trichoderma-Genus | - | - | - | 5.29 -02 | 7.23 -02 | **2.01E-02** | **6.26E-03** | **5.33E-09** | **1.12E-11** | **1.73E-11** |
| T.hara ium-pecies | - | - | - | - | 9.63 -01 | 5.21 -01 | 3.35 -01 | **2.04E-10** | **8.00E-16** | **2.65E-14** |
| Pusillae-ection | - | - | - | - | - | 6.13 -01 | 4.96 -01 | **3.34E-06** | **3.91E-09** | **7.56E-09** |
| Claviceps-Genus | - | - | - | - | - | - | 9.56 -01 | **2.98E-06** | **8.88E-10** | **2.95E-09** |
| .oxysporum-pecies | - | - | - | - | - | - | - | **3.42E-30** | **3.32E-53** | **7.78E-31** |
| Claviceps-ection | - | - | - | - | - | - | - | - | **2.05E-11** | **1.35E-04** |
| C.purpurea-pecies | - | - | - | - | - | - | - | - | - | 6.25 -01 |
| .fu i uroi-pecies | - | - | - | - | - | - | - | - | - | - |

**Table S5:** Number of duplicated genes and unique gene pairs with a pairwise identity ≥ 80% and the proportion of these gene pairs that are located next to each other (separated by 0 genes) and separated by five or fewer genes (≤ 5 genes) for all 53 *Claviceps* genomes.

| | | | | Separation | |
|---|---|---|---|---|---|
| **Species** | **Strain** | **Gene pairs† (#)** | **Duplicated genes (#)** | **0 genes** | **≤ 5 genes** |
| *C. purpurea* | 20.1 | 997 | 846 | 11.74% | 30.19% |
| *C. purpurea* | Clav04 | 578 | 710 | 8.65% | 11.94% |
| *C. purpurea* | Clav26 | 429 | 587 | 17.48% | 30.77% |
| *C. purpurea* | Clav46 | 415 | 553 | 18.55% | 29.64% |
| *C. purpurea* | Clav55 | 373 | 523 | 16.09% | 26.81% |
| *C. purpurea* | LM4 | 426 | 591 | 19.95% | 34.51% |
| *C. purpurea* | LM5 | 412 | 536 | 15.78% | 30.83% |
| *C. purpurea* | LM14 | 352 | 493 | 19.6% | 32.95% |
| *C. purpurea* | LM28 | 404 | 542 | 14.36% | 25.99% |
| *C. purpurea* | LM30 | 352 | 511 | 21.88% | 38.35% |
| *C. purpurea* | LM33 | 395 | 528 | 18.23% | 32.66% |
| *C. purpurea* | LM39 | 393 | 521 | 17.3% | 28.24% |
| *C. purpurea* | LM46 | 383 | 550 | 15.4% | 29.24% |
| *C. purpurea* | LM60 | 374 | 519 | 21.39% | 34.22% |
| *C. purpurea* | LM71 | 332 | 484 | 17.77% | 31.02% |
| *C. purpurea* | LM207 | 354 | 515 | 17.8% | 27.68% |
| *C. purpurea* | LM223 | 348 | 487 | 21.84% | 34.2% |
| *C. purpurea* | LM232 | 424 | 542 | 15.33% | 26.42% |
| *C. purpurea* | LM233 | 673 | 616 | 11.59% | 24.37% |
| *C. purpurea* | LM461 | 401 | 557 | 14.96% | 27.93% |
| *C. purpurea* | LM469 | 361 | 489 | 20.5% | 32.96% |
| *C. purpurea* | LM470 | 410 | 410 | 16.34% | 27.8% |
| *C. purpurea* | LM474 | 319 | 496 | 18.81% | 30.72% |
| *C. purpurea* | LM582 | 386 | 512 | 13.99% | 24.09% |
| *C. aff. purpurea* | Clav52 | 235 | 355 | 20.0% | 31.06% |
| *C. capensis* | CCC1504 | 144 | 247 | 13.89% | 21.53% |
| *C. pazoutovae* | CCC1485 | 182 | 270 | 14.29% | 20.33% |
| *C. monticola* | CCC1483 | 174 | 272 | 13.22% | 21.84% |
| *C. occidentalis* | LM78 | 173 | 278 | 18.5% | 26.59% |
| *C. occidentalis* | LM77 | 151 | 250 | 17.88% | 28.48% |
| *C. occidentalis* | LM84 | 431 | 313 | 10.9% | 18.79% |
| *C. quebecensis* | LM458 | 176 | 259 | 19.32% | 26.14% |
| *C. quebecensis* | Clav32 | 189 | 284 | 14.29% | 24.34% |
| *C. quebecensis* | Clav50 | 161 | 258 | 14.91% | 26.09% |
| *C. ripicola* | LM218 | 386 | 523 | 16.84% | 31.61% |
| *C. ripicola* | LM219 | 393 | 490 | 15.78% | 28.5% |
| *C. ripicola* | LM220 | 412 | 412 | 16.02% | 28.64% |
| *C. ripicola* | LM454 | 434 | 546 | 13.13% | 21.43% |
| *C. spartinae* | CCC535 | 251 | 368 | 10.36% | 16.33% |
| *C. arundinis* | CCC1102 | 431 | 518 | 11.6% | 22.51% |
| *C. arundinis* | LM583 | 362 | 442 | 15.19% | 26.24% |
| *C. humidiphila* | LM576 | 401 | 538 | 14.96% | 23.44% |
| *C. perihumidiphila* | LM81 | 351 | 494 | 19.66% | 33.62% |
| *C. cyperi* | CCC1219 | 193 | 244 | 5.7% | 7.77% |
| *C. pusilla* | CCC602 | 9 | 17 | 0.0% | 0.0% |
| *C. fusiformis* | PRL 1980 | 4 | 8 | 0.0% | 0.0% |
| *C. lovelessii* | CCC647 | 7 | 14 | 0.0% | 0.0% |
| *C. digitariae* | CCC659 | 10 | 18 | 0.0% | 0.0% |
| *C. maximensis* | CCC398 | 3 | 6 | 0.0% | 0.0% |
| *C. sorghi* | CCC632 | 12 | 23 | 0.0% | 8.33% |
| *C. africana* | CCC489 | 8 | 16 | 0.0% | 0.0% |
| *C. citrina* | CCC265 | 24 | 34 | 4.17% | 4.17% |
| *C. paspali* | RRC 1481 | 1 | 2 | 0.0% | 0.0% |

† Unique pairs (i.e. pairs of gene A : gene B and gene B : gene A are not counted twice).

**Table S6:** Means, standard deviations, and additional statistics of repeat-induced point mutation (RIP) composite indexes and large RIP affected regions (LRARs) for all 53 *Clvaiceps* genomes computed using The RIPper on default settings.

| Species | Strain | Section | RIP composite index† | RIP affected windows (#) | RIP genomic content (%) | LRARs (#) | LRARs length (kbp) | LRARs genomic content (kbp) | LRARs composite index† | LRARs GC content (%) |
|---|---|---|---|---|---|---|---|---|---|---|
| *C. purpurea* | 20.1 | Claviceps | -0.61 ± 0.29 | 80 | 0.12% | | | | | |
| *C. purpurea* | Clav04 | Claviceps | -0.59 ± 0.28 | 136 | 0.21% | | | | | |
| *C. purpurea* | Clav26 | Claviceps | -0.59 ± 0.28 | 86 | 0.14% | | | | | |
| *C. purpurea* | Clav46 | Claviceps | -0.59 ± 0.28 | 88 | 0.14% | | | | | |
| *C. purpurea* | Clav55 | Claviceps | -0.59 ± 0.28 | 96 | 0.15% | | | | | |
| *C. purpurea* | LM4 | Claviceps | -0.59 ± 0.28 | 75 | 0.12% | | | | | |
| *C. purpurea* | LM5 | Claviceps | -0.59 ± 0.28 | 75 | 0.12% | | | | | |
| *C. purpurea* | LM14 | Claviceps | -0.59 ± 0.28 | 66 | 0.11% | | | | | |
| *C. purpurea* | LM28 | Claviceps | -0.59 ± 0.28 | 79 | 0.13% | | | | | |
| *C. purpurea* | LM30 | Claviceps | -0.59 ± 0.28 | 71 | 0.12% | | | | | |
| *C. purpurea* | LM33 | Claviceps | -0.59 ± 0.28 | 76 | 0.12% | | | | | |
| *C. purpurea* | LM39 | Claviceps | -0.59 ± 0.28 | 68 | 0.11% | | | | | |
| *C. purpurea* | LM46 | Claviceps | -0.59 ± 0.28 | 84 | 0.14% | | | | | |
| *C. purpurea* | LM60 | Claviceps | -0.59 ± 0.28 | 61 | 0.1% | | | | | |
| *C. purpurea* | LM71 | Claviceps | -0.59 ± 0.28 | 79 | 0.13% | | | | | |
| *C. purpurea* | LM207 | Claviceps | -0.59 ± 0.28 | 78 | 0.13% | | | | | |
| *C. purpurea* | LM223 | Claviceps | -0.59 ± 0.28 | 94 | 0.15% | | | | | |
| *C. purpurea* | LM232 | Claviceps | -0.59 ± 0.28 | 64 | 0.1% | | | | | |
| *C. purpurea* | LM233 | Claviceps | -0.59 ± 0.28 | 73 | 0.12% | | | | | |
| *C. purpurea* | LM461 | Claviceps | -0.59 ± 0.28 | 72 | 0.12% | | | | | |
| *C. purpurea* | LM469 | Claviceps | -0.59 ± 0.28 | 76 | 0.12% | | | | | |
| *C. purpurea* | LM470 | Claviceps | -0.59 ± 0.28 | 90 | 0.15% | | | | | |
| *C. purpurea* | LM474 | Claviceps | -0.59 ± 0.28 | 94 | 0.15% | | | | | |
| *C. purpurea* | LM582 | Claviceps | -0.59 ± 0.28 | 81 | 0.13% | | | | | |
| *C. aff. purpurea* | Clav52 | Claviceps | -0.59 ± 0.31 | 65 | 0.11% | | | | | |
| *C. capensis* | CCC1504 | Claviceps | -0.61 ± 0.27 | 41 | 0.07% | | | | | |
| *C. pazoutovae* | CCC1485 | Claviceps | -0.61 ± 0.27 | 39 | 0.07% | | | | | |
| *C. monticola* | CCC1483 | Claviceps | -0.59 ± 0.28 | 45 | 0.08% | | | | | |
| *C. occidentalis* | LM78 | Claviceps | -0.55 ± 0.31 | 119 | 0.2% | | | | | |
| *C. occidentalis* | LM77 | Claviceps | -0.55 ± 0.31 | 133 | 0.23% | | | | | |
| *C. occidentalis* | LM84 | Claviceps | -0.55 ± 0.31 | 111 | 0.19% | | | | | |
| *C. quebecensis* | LM458 | Claviceps | -0.57 ± 0.29 | 88 | 0.15% | | | | | |
| *C. quebecensis* | Clav32 | Claviceps | -0.57 ± 0.29 | 83 | 0.14% | 2 | 4.5 ± 0.01 | 9.0 | 0.73 ± 0.05 | 54.07% ± 0.37% |
| *C. quebecensis* | Clav50 | Claviceps | -0.57 ± 0.29 | 76 | 0.13% | | | | | |
| *C. ripicola* | LM218 | Claviceps | -0.57 ± 0.29 | 82 | 0.13% | | | | | |
| *C. ripicola* | LM219 | Claviceps | -0.57 ± 0.28 | 87 | 0.14% | | | | | |
| *C. ripicola* | LM220 | Claviceps | -0.58 ± 0.29 | 85 | 0.14% | | | | | |
| *C. ripicola* | LM454 | Claviceps | -0.57 ± 0.28 | 82 | 0.13% | | | | | |
| *C. spartinae* | CCC535 | Claviceps | -0.58 ± 0.28 | 73 | 0.12% | | | | | |
| *C. arundinis* | CCC1102 | Claviceps | -0.58 ± 0.28 | 55 | 0.09% | | | | | |
| *C. arundinis* | LM583 | Claviceps | -0.58 ± 0.28 | 60 | 0.1% | | | | | |
| *C. humidiphila* | LM576 | Claviceps | -0.59 ± 0.28 | 64 | 0.1% | | | | | |
| *C. perihumidiphila* | LM81 | Claviceps | -0.58 ± 0.28 | 80 | 0.13% | | | | | |
| *C. cyperi* | CCC1219 | Claviceps | -0.60 ± 0.27 | 90 | 0.17% | 1 | 4.5 ± 0 | 4.5 | 0.66 ± 0.00 | 53.18% ± 0.00% |
| *C. pusilla* | CCC602 | Pusillae | 0.15 ± 1.04 | 36,205 | 38.36% | 564 | 13.7 ±11.2 | 7,739 | 1.30 ± 0.17 | 25.23% ± 4.05% |
| *C. fusiformis* | PRL 1980 | Pusillae | 0.03 ± 1.30 | 18,107 | 16.67% | 274 | 5.9 ± 1.6 | 1,610 | 1.84 ± 0.47 | 4.81% ± 3.15% |
| *C. lovelessii* | CCC647 | Pusillae | -0.02 ± 1.00 | 25,695 | 30.29% | 399 | 10.8 ± 6.6 | 4,320 | 1.32 ± 0.16 | 23.55% ± 3.80% |
| *C. digitariae* | CCC659 | Pusillae | -0.25 ± 0.89 | 13,661 | 20.17% | 271 | 11.5 ± 7.5 | 3,109 | 1.37 ± 0.15 | 22.13% ± 3.38% |
| *C. maximensis* | CCC398 | Pusillae | -0.24 ± 0.91 | 13,517 | 20.37% | 148 | 14.6 ± 13.5 | 2,156 | 1.43 ± 0.13 | 21.44% ± 2.49% |
| *C. sorghi* | CCC632 | Pusillae | 0.01 ± 1.04 | 21,622 | 29.57% | 348 | 13.8 ± 13.4 | 4,804 | 1.41 ± 0.16 | 23.68% ± 2.79% |
| *C. africana* | CCC489 | Pusillae | 0.04 ± 0.99 | 25,266 | 33.09% | 289 | 22.0 ± 19.7 | 6,362 | 1.33 ± 0.09 | 20.11% ± 3.85% |
| *C. citrina* | CCC265 | Citrinae | 0.36 ± 1.11 | 43,520 | 48.69% | 503 | 9.9 ± 6.4 | 4,957 | 1.34 ± 0.17 | 29.80% ± 3.95% |
| *C. paspali* | RRC 1481 | Paspalorum | -0.35 ± 1.00 | 5,351 | 9.05% | 131 | 6.1 ± 1.6 | 799 | 1.85 ± 0.39 | 4.11% ± 2.75% |

† *Composite Index Value* [(TpA/ ApT) – (CpA + TpG/ ApC + GpT)], positive values imply RIP
